# Divergent Recruitment of Developmentally-Defined Neuronal Ensembles Supports Memory Dynamics

**DOI:** 10.1101/2023.11.06.565779

**Authors:** Vilde A. Kveim, Laurenz Salm, Talia Ulmer, Steffen Kandler, Fabia Imhof, Flavio Donato

**Author notes:** Corresponding author: Flavio Donato.

## Abstract

Memories are dynamic constructs whose properties change with time and experience. The biological mechanisms underpinning these dynamics remain elusive, particularly concerning how shifts in the composition of memory-encoding neuronal ensembles influence a memory properties’ evolution over time. By leveraging a developmental approach to target distinct subpopulations of principal neurons, we show that memory encoding results in the concurrent establishment of multiple memory traces in the mouse hippocampus. Two of these traces are instantiated in subpopulations of early- and late-born neurons and follow distinct reactivation trajectories post-encoding. Notably, the divergent recruitment of these subpopulations underpins memory ensembles’ gradual reorganization, and modulates memory persistence and plasticity across multiple learning episodes. Thus, our findings reveal profound and intricate relationships between ensemble dynamics and memories’ progression over time.

---

The ability to create memories that persist through time is crucial for survival, as it allows individuals to learn from past experiences and select appropriate behaviours when similar contingencies are encountered again (1–3). In mammals, this ability is critically supported by a network of interconnected areas centred around the hippocampus (3–7). In the hippocampus, subsets of neurons that during an experience are characterized by enhanced excitability, synaptic plasticity, and the expression of immediate early genes (IEGs) such as c-Fos, function as correlated neuronal ensembles to become part of the biological trace, or Engram, where the memory of that event is encoded (8–18).

Classically, memory persistence has been attributed to the capacity to reinstate at later times patterns of neuronal activity that were initially established during memory acquisition; a phenomenon commonly referred to as memory recall (9, 10, 19). In line with this idea, activity patterns that in humans are observed in specific brain areas during the encoding of a memory are similar to those observed during its retrieval (20–22). Similarly, the c-Fos+ neuronal ensemble that in rodents is established during a memory’s acquisition overlaps with the one established during its recall, with the extent of this overlap being associated with retrieval success (9, 10, 14, 16, 23–26). Furthermore, the reactivation of the c-Fos+ ensemble established at acquisition is both necessary for memory retrieval in physiological conditions, and sufficient to induce artificial recollection of a specific memory at later times (11, 13, 14, 24, 27). Together, these data have supported a reactivation-centred model, according to which memory encoding leads to the establishment of a distributed network of memory ensembles in multiple cortical and subcortical structures, with hippocampal ensembles serving as an index for the reactivation of this memory network during recall (8, 10, 12, 17, 28–30).

Recent evidence has additionally shed light on the dynamic nature of memories, revealing that memories behave as dynamic constructs that change over time and with experience. Their properties, such as specificity, accessibility, or susceptibility to disruption, evolve after encoding, and their content can be updated upon changes in external or internal contingencies (*25, 31–42*). Likewise, activity patterns in brain areas where memory ensembles are established upon learning, like the hippocampus, evolve over time and experience (*2, 14, 27, 43–49*). New sets of neurons are recruited into memory ensembles over multiple learning sessions, while a neuron’s probability of reactivation upon successive exposures to the same experience decreases as a function of time elapsed between them (a phenomenon known as “representational drift”) (*43, 50–57*). These discoveries on the dynamic nature of memories and ensemble activity have raised several fundamental questions. What is the underlying logic that governs the reorganization of memory ensembles linked to a specific memory? How do dynamic changes in supporting ensembles affect a memory’s persistence over time? And finally, are hippocampal ensembles recruited at different stages of a memory’s lifetime always composed of similar neurons? Or alternatively, are distinct neuronal subpopulations recruited at specific times to confer dynamic properties to a memory?

Providing an answer to these questions is crucial to understanding how the hippocampus participates in memory processes. It has now been established that hippocampal principal neurons segregate into distinct subpopulations along a series of variables, from gene expression to anatomical features, local and long-range connectivity, maturation trajectories, intrinsic properties, functional tuning, and structural and functional plasticity upon learning (*58–70*). Strikingly, the logic by which this diversity is organized in hippocampal circuits is inherently tied to development, since populations of neurons born on the same embryonic day (referred to as “isochronic neurons”) exhibit similar properties and might function as preconfigured functional units upon experience, while neurons belonging to different isochronic cohorts segregate into distinct functional subpopulations (*58, 59, 66, 71*). Therefore, one major question is whether the composition of hippocampal ensembles supporting the processing of a specific memory evolves along specific trajectories, anchored in distinct developmentally-defined neuronal subpopulations. Such subpopulations could participate differentially in memory processes at specific stages of a memory’s lifetime, leading to a dynamic reorganization in ensemble composition that might parallel, and potentially facilitate, changes in a memory’s properties, underpinning memory dynamics over time.

In this study, we set out to directly test this hypothesis by focusing specifically on developmentally stratified hippocampal principal neurons, and by dissecting their roles in memory processes unfolding in adult male and female mice. We examined memory ensembles established in the CA3 subfield of the hippocampus, as this area exhibits key features of an associative memory network and plays a crucial role in memory processing throughout a memory’s lifespan (*14, 16, 46, 72*). We particularly emphasized the processing of fear-associated, hippocampus-dependent memories, as their encoding is time-constrained and the temporal evolution of their properties follows a consistent pattern (*33, 34, 36*). Our experiments reveal that in the hippocampal network, learning results in the parallel establishment of two distinct memory traces that exhibit unique trajectories for their emergence, reactivation, and permanence. These traces are instantiated in distinct neurogenesis-defined subpopulations of early- and late-born neurons, whose timely activation during memory acquisition, consolidation, or recall underpins a memory’s long-term persistence and availability to retrieval. Last, we show that divergent recruitment of early- and late-born neurons to memory ensembles in the days after encoding modulates the temporal boundaries of a transient window when the integration of information across multiple learning episodes into cohesive memories is facilitated. Together, our experiments reveal that ensemble dynamics, anchored in the divergent recruitment of developmentally-defined hippocampal memory traces established at acquisition, underpin the encoding, persistence, and evolution of a memory’s properties over time.

## Early- and Late-Born Neurons exhibit distinct recruitment dynamics to parallel-emerging memory traces

To dissect how developmentally-defined subpopulations of hippocampal principal neurons contribute to memory ensembles supporting the encoding and retrieval of fear-associated memories, we used a multi-step approach. First, we employed a pulse-chase method to label neurons with similar birthdates; next, we used an IEGs-based method to identify c-Fos-expressing (c-Fos+) neuronal ensembles recruited at various stages of a contextual Fear Conditioning (cFC) memory’s lifetime (Fig. 1A). Therefore, pregnant dams were injected with 5-bromo-2′-deoxyuridine (BrdU) at specific times during gestation (*i.e.* on different Embryonic days, E-day), to target CA3 neurons born around the time of injection (Fig. 1, A and B). After reaching adulthood (*i.e.* older than 60 post-natal days, >P60), distinct cohorts of BrdU-labelled mice were trained in the cFC task. In this task, the memory for the association between a conditioning stimulus (CS, the Training Context) and an aversive stimulus (US, a foot shock) is rapidly encoded during a brief session (referred to as “Acquisition” session from now on, Fig. 1A). The retrieval of the fear memory faithfully occurs upon the presentation of the Training Context even at significant temporal delays from encoding (referred to as “Recall” session from now on, Fig. 1A), and induces an increase in both the amount of time mice spend freezing in the Training Context, as well as in the prevalence of c-Fos+ neurons in hippocampal subfield CA3 (“c-Fos+ ensemble”, Fig. S1, A-C) (*14, 16, 46*). In our experiments, the expression of c-Fos in specific isochronic subpopulations was systematically mapped by quantifying the fraction of double-positive BrdU+, c-Fos+ neurons over the entire isochronic cohort (BrdU+, Fig. 1C). Through the course of our study, we will focus on two populations of isochronic neurons, those born on either E12 (from now on defined as “Early Born Neurons”, or EBNs) or on E15 (“Late Born Neurons”, LBNs, Fig. 1B), since previous studies have shown that these subpopulations are maximally distinct in terms of intrinsic properties, activity dynamics, and connectivity (*58, 67, 71*).

**Fig. 1.**
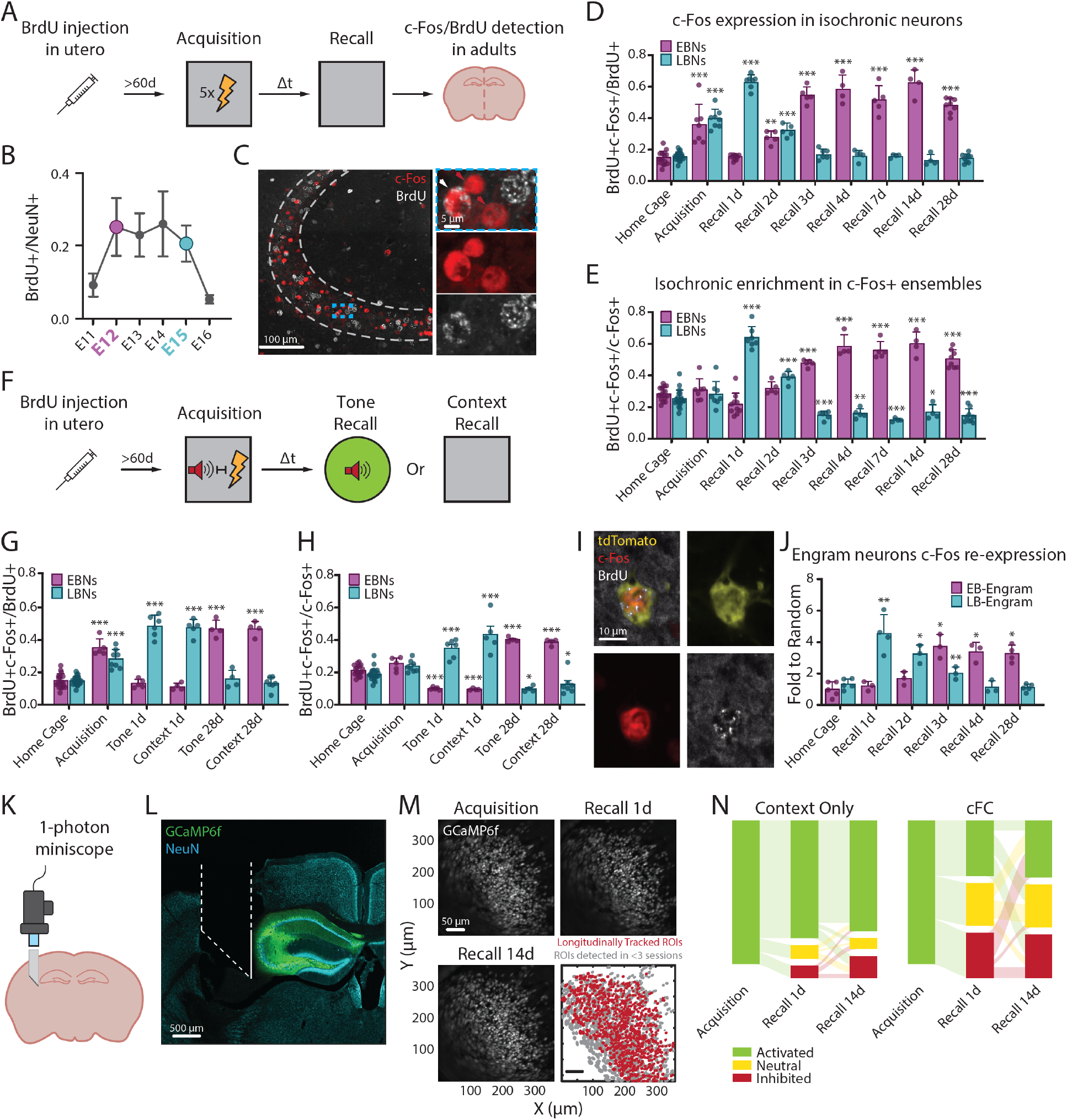
Divergent recruitment of EBNs and LBNs to memory ensembles throughout a memory’s lifetime. **(A)** Schematic of the pulse-chase labelling protocol in association with contextual Fear Conditioning (cFC) learning task. The Recall session is performed at a variable delay Δt from the Acquisition session (1 day ≤ Δt ≤ 28 days). **(B)** Fraction of CA3 (NeuN+) neurons positive for BrdU (y-axis) plotted as a function of the day of BrdU injection during embryonic development (x-axis). Each data point is the mean ± SD calculated over at least 3 animals. Early Born Neurons (EBNs, magenta) are labelled by the injection of BrdU on E12, and Late Born Neurons (LBNs, cyan) on E15. **(C)** c-Fos (red) and BrdU (white) immunolabelling of neurons in the pyramidal layer (dashed line) of hippocampal area CA3. The inset area (dashed blue box) is magnified on the right, for which individual images for c-Fos (red) and BrdU (white) are shown. The red arrow points to a c-Fos+, BrdU-neuron, white arrow points to a c-Fos+, BrdU+ neuron. Images are maximum intensity projections, 30X silicone objective. **(D)** Fraction of CA3 EBNs (magenta) or LBNs (cyan) expressing c-Fos (BrdU+ c-Fos+/BrdU+) upon the encoding and retrieval of a cFC memory. Each dot is a different mouse. Bars represent the mean, error bars the SD. EBNs exhibited a significant increase in c-Fos expression upon cFC Acquisition and Recall, only when this happened at least 2 days after Acquisition (one-way ANOVA, F(8, 55) = 58.18, p<0.0001. Dunnett’s multiple comparisons test to Home Cage EBNs: p<0.0001 for Acquisition and Recall ≥3d; p = 0.003 for Recall 2d; p>0.05 for Recall 1d). LBNs exhibited a significant increase in c-Fos expression upon cFC Acquisition and Recall, only when this happened less than 3 days after Acquisition (one-way ANOVA, F(8, 60) = 161.8, p<0.0001. Dunnett’s multiple comparisons test to Home Cage LBNs: p<0.0001 for Acquisition and Recall 1 and 2 days; p>0.05 for Recall ≥3d). At least 4 animals per group, no repeated measures. **(E)** Prevalence of EBNs or LBNs in CA3 c-Fos+ ensembles (BrdU+ c-Fos+/c-Fos+) supporting the encoding and retrieval of a cFC memory. Each dot is a different mouse. Bars represent the mean, error bars the SD. EBNs abundance in ensembles established at Acquisition and Recall 1d and 2d was comparable to baseline levels, but significantly higher if Recall was performed thereafter (one-way ANOVA, F(8, 55) = 48.38, p<0.0001. Dunnett’s multiple comparisons test to Home Cage: p>0.05 for Acquisition, Recall 1d and 2d; p<0.0001 for Recall ≥3d). Conversely, LBNs abundance in ensembles established at Acquisition was comparable to baseline levels, higher for Recall performed 1d or 2d after, but significantly lower thereafter (one-way ANOVA, F(8, 59) = 64.69, p<0.0001. Dunnett’s multiple comparisons test to Home Cage LBNs: p>0.05 for Acquisition; p<0.0001 for Recall 1d and 2d; p = 0.0006 for Recall 3d; p = 0.0059 for Recall 4d; p = 0.0002 for Recall 7d; p = 0.032 for Recall 14d; p<0.0001 for Recall 28d). At least 4 animals per group, no repeated measures. **(F)** Schematic of the “trace Fear Conditioning” (tFC) protocol on the pulse-chased mice. In this task, a specific tone (Tone-CS) and the delivery of a foot shock (the US) are presented in a specific context (Context-CS) and separated by a temporal delay (*i.e.* a trace) of several seconds; hippocampal activity is necessary for the integration of information across time and the encoding of an association between the Tone-CS and the US. Upon tFC Recall, memory retrieval can be induced by either reintroducing conditioned animals into the Training Context (“Context Recall”), or by presenting the Tone-CS in a novel, neutral context (“Tone Recall”). **(G)** Fraction of CA3 EBNs (magenta) or LBNs (cyan) expressing c-Fos (BrdU+ c-Fos+/BrdU+) upon the encoding and retrieval of a tFC memory. Each dot is a different mouse. Bars represent the mean, error bars the SD. EBNs exhibited a significant increase in c-Fos expression upon tFC Acquisition, and when the fear memory was retrieved 28, but not 1, days after encoding by either the presentation of the Tone- or Context-CS (one-way ANOVA, F(5,35) = 83.23, p<0.0001. Dunnett’s multiple comparisons test to Home Cage c-Fos expression in EBNs: p<0.0001 for Acquisition and both Context and Tone recall at 28d; p>0.05 for Tone and Context Recall at 1 day). LBNs exhibited a significant increase in c-Fos expression upon tFC Acquisition, and when the fear memory was retrieved 1, but not 28, days after encoding by either the presentation of the Tone- or Context-CS (one-way ANOVA, F(5,47) = 100.5, p<0.0001. Dunnett’s multiple comparisons test to Home Cage c-Fos expression in LBNs: p<0.0001 for Acquisition and both Context and Tone recall at 1d; p>0.05 for Tone and Context Recall at 28 days). At least 4 animals per group, no repeated measures. **(H)** Prevalence of EBNs or LBNs in CA3 c-Fos+ ensembles (BrdU+ c-Fos+/c-Fos+) supporting the encoding and retrieval of a tFC memory. Each dot is a different mouse. Bars represent the mean, error bars the SD. EBNs abundance in c-Fos+ ensembles established at Acquisition was comparable to baseline, significantly lower for both Tone and Context Recall performed at 1d, and significantly higher for both Tone and Context Recall performed at 28d (one-way ANOVA, F(5,35) = 100.5, p<0.0001. Dunnett’s multiple comparisons test to Home Cage: p<0.0001 for both Context and Tone recall either at 1d and 28d; p>0.05 for Acquisition). LBNs abundance in c-Fos+ ensembles established at Acquisition was comparable to baseline, significantly higher for both Tone and Context Recall performed at 1d, and significantly lower for both Tone and Context Recall performed at 28d (one-way ANOVA, F(5,46) = 33.42, p<0.0001. Dunnett’s multiple comparisons test to Home Cage: p<0.0001 for both Context and Tone recall at 1d; p = 0.0132 for Tone Recall 28d; p = 0.0368 for Context Recall 28d; p>0.05 for Acquisition). At least 4 animals per group, no repeated measures. **(I)** tdTomato (yellow), c-Fos (red), and BrdU (white) immunolabelling of LB-Engram neurons. Images are maximum intensity projections, 30X silicone objective. **(J)** Rate of c-Fos Re-expression in Early Born (EB-) and Late Born (LB-) CA3 Engram neurons at Recall. Each dot is a different mouse. Bars represent the mean, error bars the SD. In each experimental group, the rate of reactivation of EB- and LB-Engram neurons was expressed as the fraction of triple positive tdTomato+, BrdU+, and c-Fos+ neurons normalized by the number of tdTomato+ neurons. These values were compared to the probability that a neuron would be positive to all markers by chance (calculated as p_(tdTomato+)_* p_(c-Fos+)_* p_(BrdU+)_ See Methods). Two-tailed one-sample t-test analysis indicates that for EB-Engram neurons, re-expression rates were at chance levels for Recall 1d (t(2) = 1.205, p>0.05) and Recall 2d (t(2) = 2.972, p>0.05), but higher than what would be expected by chance upon Recall 3d (t(2) = 8.807, p = 0.015), Recall 4d (t(2) = 6.691, p = 0.0216), and Recall 28d (t(2) = 5.943, p = 0.027). Conversely, for LB-Engram neurons, re-expression rates were significantly higher than chance upon Recall 1d (t(3) = 5.857, p = 0.01), Recall 2d (t(2) = 6.673, p = 0.022), and Recall 3d (t(3) = 18.37, p = 0.003), but they were at chance levels for Recall 4d (t(2) = 0.523, p>0.05) and Recall 28d (t(4) = 0.866, p>0.05). One-way ANOVA identifies significant differences in the rate of re-expression of EB-Engram neurons (F(5,15) = 14.93, p<0.0001. Dunnett’s multiple comparisons test to Home Cage: p>0.05 for Recall 1d and 2d; p<0.0001, p = 0.0003, and p = 0.0002 for Recall 3d, 4d, and 28d, respectively) and LB-Engram neurons (F(5,17) = 20.99, p<0.0001. Dunnett’s multiple comparisons test to Home Cage: p<0.0001 and p = 0.0022 for Recall 1d and 2d, respectively; p>0.05 for Recall 3d, 4d, and 28d). At least 3 animals per group, no repeated measures. **(K)** Schematic representation of the implant location for recording the activity of large populations of CA3 neurons in freely moving animals. **(L)** Representative histology revealing the position of the endomicroscope (white dashed outline) and the prism imaging face (solid white outline) in relation to the hippocampal formation. GCaMP6f expressing neurons in green, and NeuN expressing neurons in light blue. The image is a single confocal plane, 4X air objective. **(M)** Longitudinal tracking of the same neurons within one recording Field of View (FOV). Activity from the same neurons could be recorded longitudinally across experimental sessions spaced up to 14 days apart. Images are Maximum Intensity Projections from the miniscope FOV. White is the GCaMP6f signal. Bottom right: distribution of the recorded Regions of Interest (ROIs) computed to identify neurons that could be reliably tracked across all sessions (red) and neurons that could only be detected in 1 or 2 sessions (dark grey). **(N)** Changes in the activity of individual neurons across sessions. Individual neurons were categorized as Activated (green), Neutral (yellow), or Inhibited (red) based on their change in firing rate between Home Cage and Training Context (see Methods). Only neurons that were classified as Activated during the Acquisition session are analysed (cFC cohort: 413 neurons over 5 animals. Context Only cohort: 183 neurons over 5 animals). The data are plotted as a Sankey diagram, where the three categories represent the nodes, and the thickness of the link between nodes represents the proportion of neurons that flow across two nodes (from left to right). A χ2 test conducted on a contingency table considering each possible path through the diagram as categorical variables (*i.e.* Activated-Activated-Activated, Activated-Neutral-Activated, etc.) revealed a significant difference between dynamics in the two cohorts: χ2(8, N = 100) = 97.12, p<0.0001.

Although upon memory encoding, the prevalence of c-Fos+ neurons was significantly higher than baseline (Home Cage levels) in both isochronic subpopulations, EBNs and LBNs exhibited distinct c-Fos expression dynamics upon memory retrieval (Fig. 1D). The prevalence of c-Fos+ neurons among LBNs was significantly elevated if Recall was performed up to 2 days after encoding, but was not different than baseline for Recall on day 3 or thereafter (Fig. 1D). In stark contrast, the prevalence of c-Fos+ neurons among EBNs was at baseline levels if Recall was performed 1 day after Acquisition, but steadily increased over time and became significantly elevated for Recalls performed by day 2 and later. We then asked if similarly, the prevalence of EBNs and LBNs in c-Fos+ ensembles followed a similar time course and changed as a function of the delay between memory Acquisition and Recall. For this, we normalized the number of double-positive BrdU+ and c-Fos+ neurons over the entire c-Fos+ population. Consistently, while the c-Fos+ ensemble established at Acquisition recruited similar proportions of EBNs and LBNs (reflecting their relative abundance in the hippocampal network), ensembles formed upon Recall were enriched in LBNs if this happened the day after Acquisition (Fig. 1E). Such bias rapidly disappeared, and from day 3 onwards, Recall-induced c-Fos+ ensembles were significantly shifted towards the preferential enrichment in EBNs (Fig. 1E). Importantly, we could not observe neither a differential expression of c-Fos among EBNs and LBNs, or differential recruitment in c-Fos+ ensembles if animals were never exposed to the US during training (“Context Only” controls, Fig. S1, D-H), or if they were never subjected to a Recall session after Acquisition (“No Recall” controls, Fig. S1, I-L), or if animals were introduced into a novel, neutral context during Recall (“Context B Recall” controls, Fig. S1, M-Q). Thus, our results reveal that EBNs and LBNs have distinct time courses of recruitment in hippocampal c-Fos+ ensembles underpinning the encoding and retrieval of a specific contextual-fear memory.

We then investigated if distinct recruitment dynamics could be observed upon the retrieval of a distinct hippocampus-dependent, fear-association memory. For this, we used the trace Fear Conditioning (tFC) task, which relies on the causal association between CS and US across temporal delay periods at Acquisition (30 seconds in our experiment). The tFC task further leverages multiple sensory modalities at Recall (*i.e.* exploits the presentation of either a vision-biased Context-CS or the auditory-biased Tone-CS to promote memory retrieval), and similarly to the cFC task, is hippocampus-dependent (See Methods, Fig. 1F and Fig. S1, R and S) (*73*). Consistently with the processing of a cFC memory, c-Fos expression in EBNs and LBNs (Fig. 1G), and the prevalence of these isochronic populations in memory ensembles (Fig. 1H), were a function of the temporal delay between Acquisition and Recall, irrespectively of the sensory modality exploited for memory retrieval. Thus, distinct recruitment dynamics of EBNs and LBNs were not contingent upon the engagement of specific sensory modalities at Recall, and could be observed across hippocampus-dependent tasks leveraging both spatial and temporal information.

Though these experiments revealed that EBNs and LBNs are differentially recruited into c-Fos+ ensembles established at Recall when this is performed at different stages of a memory’s lifetime, they fell short of revealing if similar dynamics could be observed in neurons that are recruited to memory ensembles upon both memory encoding and retrieval, *i.e.* in Engram neurons that are allocated to memory ensembles during Acquisition and re-express c-Fos during Recall (*8*). Hence, to investigate if Engram neurons exhibit distinct c-Fos re-expression dynamics as a function of their birthdate, we implemented an intersectional strategy. First, we injected BrdU to target EBNs and LBNs in a transgenic line that leverages the c-Fos dependent expression of the inducible recombinase CreERT2, in combination with the Cre-dependent expression of a fluorescent marker, to permanently label Engram neurons (TRAP2/Floxed-tdTomato mouse line) (*47*). Hence, when BrdU-injected animals reached adulthood, we trained them in the cFC paradigm and injected 4-Hydroxytamoxifen immediately after memory Acquisition, to trigger Cre-driven recombination and reveal Early- or Late-Born Engram neurons (EB-Engram and LB-Engram neurons, respectively) by their co-localization of BrdU and tdTomato (Fig. 1I). We then reintroduced BrdU, tdTomato labelled animals into the Training Context for memory Recall, and quantified the probability of EB- and LB-Engram neurons’ c-Fos re-expression against the values expected by chance (see Methods). Upon Recall, the fraction of LB-Engram neurons expressing c-Fos was significantly higher than chance when the cFC memory was retrieved up to 3 days after encoding, but fell to chance levels for a Recall on day 4 or 28 (Fig. 1J). Conversely, c-Fos expression in EB-Engram neurons was at chance levels for memory retrieval up to 2 days after encoding, but rose to be significantly higher than chance for Recall on days 3, 4, or 28 after acquisition (Fig. 1J).

These findings suggest that neurons active during the Acquisition of a cFC memory might segregate into distinct subpopulations, each with unique reactivation patterns during Recall. One group of neurons would be reactivated at Recall in the days immediately following Acquisition but not after a longer delay, reflecting c-Fos expression dynamics of LB-Engram neurons. In contrast, another group might be reactivated mainly when Recall happens after a longer temporal gap, similar to c-Fos expression dynamics of EB-Engram neurons. We, therefore, decided to use an orthogonal approach to directly monitor the activity of a large population of CA3 neurons longitudinally, across three experimental sessions: Acquisition, Recall after 1 day, and Recall after 14 days. To this end, we used a head-mounted miniaturized 1-photon microscope (miniscope) to record CA3 network activity through calcium imaging with single-cell resolution in freely moving mice (Fig. 1, K-M). Calcium imaging was performed on two distinct cohorts of animals that were trained in either the cFC or Context Only tasks (Fig. S2, A and B). For the cFC cohort, freezing levels were elevated at any time after Acquisition, indicating that the repeated Recall did not lead to a quantifiable extinction of the fear memory (Fig. S3A). The Context Only cohort instead exhibited minimal levels of freezing at any moment during the experiment (Fig. S3A). On each experimental session – designated as “Acquisition”, “Recall 1d”, and “Recall 14d” – we first recorded CA3 network activity while animals were in their Home Cage, to infer baseline levels of activity for each neuron. We then introduced each mouse in the Training Context, to record activity from the same neurons during memory Acquisition or Recall (Fig. S3, B and C). Longitudinally tracked neurons were classified as “Activated”, “Neutral”, or “Inhibited” based on the difference in activity rate exhibited in the Training Context vs. Home Cage imaging periods (see Methods).

Among neurons that were Activated at Acquisition (*i.e.* putative engram neurons), and in addition to a group of neurons that were consistently Activated across the three experimental sessions, a large fraction showed reactivation dynamics in line with our LB- and EB-Engram labelling experiments. One group of neurons was reactivated on Recall 1d but not on Recall 14d, thereby showing activity dynamics that recapitulated c-Fos expression in LB-Engram neurons (Fig. 1N). In stark contrast, another group were reactivated on Recall 14d but not Recall 1d, thereby showing activity dynamics that recapitulated c-Fos expression in EB-Engram neurons. Moreover, the proportion of neurons exhibiting LB- or EB-Engram like reactivation dynamics was significantly higher in animals trained in the cFC task than in the cohort that was trained in the Context Only task, where neurons followed reactivation dynamics that were largely consistent with previous reports of representational drift in the hippocampus (Fig. 1N) (*52, 55*). Thus, the dynamic reorganization of active neuronal ensemble observed in the longitudinal recording experiments was in line with results obtained with the Engram labelling technique, and consistent with a more pronounced reorganization of memory ensembles established upon Recall of a cFC memory when compared to the memory of a neutral context (as suggested by the isochronic recruitment experiments in Fig. S1, G and H).

These results additionally implied that upon the processing of a cFC memory, a significant fraction of neurons changed their responses to the Training Context between Recall 1d and Recall 14d. Thus, for every longitudinally tracked neuron on each experimental session, we calculated a Tuning Score (TS) expressing the measure in which a neuron’s activity rate would increase (TS>0) or decrease (TS<0) during the exploration of the Training Context in comparison to the Home Cage (see Methods) (74). We then assessed the stability of single neurons’ tuning over time by plotting TS values across each session’s pairing. For the Context Only cohort, most neurons maintained consistent tuning across all pairs (“Acquisition vs. Recall 1d”, “Acquisition vs. Recall 14d”, and “Recall 1d vs. Recall 14d”) (Fig. S3D, top row). Kernel Density Estimate (KDE) analysis confirmed this, indicating a high density of data points along the diagonal (Fig. S3E, top row). For the cFC cohort, similar distributions emerged when comparing Acquisition with either Recall session (Fig. S3, D and E bottom row, left and centre). However, a distinct pattern arose when comparing the two cFC Recall sessions against each other. Neurons displayed a broader distribution, with many found off-diagonal, suggesting numerous neurons changed their response to the Training Context between the two (Fig. S3, D and E bottom row, right). To quantify this phenomenon, we calculated the difference in each neuron’s Tuning Scores (ΔTS) across session pairs, and analysed the cumulative distribution of ΔTS across the recorded population. As expected, the Context Only cohort showed consistent cumulative distributions for all session pair comparisons (Fig. S3F, left). In contrast, the distribution for the comparison between 1d and 14d Recall of a cFC memory significantly diverged from those comparing each cFC Recall to Acquisition (Fig. S3F, right). Thus, a higher proportion of cells changed their responses to the Training Context between 1 and 14 days after Acquisition in cFC-trained animals than in the Context Only cohort. This confirms distinct reactivation dynamics in neuronal ensembles supporting 1d and 14d Recall of a cFC memory.

Taking these results together, we advanced the hypothesis that upon the encoding of a hippocampus-dependent fear memory, two distinct memory traces are laid down in the CA3 network. A transient, fast-maturing but short-lived memory trace, instantiated in a subpopulation of late-born neurons, would be preferentially reactivated during memory Recall the day after Acquisition, with its reactivation rate rapidly decaying over the following days. At the same time, an enduring, slow-maturing and long-lasting memory trace, instantiated in a subpopulation of early-born neurons, would be encoded within the network, with its reactivation rates at Recall steadily increasing within days after encoding, and remaining sustained at remote time points. The shift in recruitment between these two traces would drive memory ensembles’ gradual reorganization, whose underlying logic is grounded in the alternative recruitment of developmentally-defined subpopulations of late- and early-born neurons as a function of time elapsed between Acquisition and Recall.

## Timely recruitment of isochronic subpopulations is necessary for memory recall

The divergent recruitment dynamics of EBNs and LBNs led us to investigate if memory recall relies on the activation of different hippocampal subpopulations when memories are retrieved at different intervals after their initial encoding. Thus, we set out to determine whether the recruitment of EBNs and LBNs to memory ensembles formed during Recall, specifically at times when c-Fos+ expression was at its peak within each subpopulation and maximally different across subpopulations (*i.e.* Recall 28d and 1d, respectively, as depicted in Fig. 1E), had a functional impact on memory retrieval. To address this question, we employed a multi-step, neurogenesis-targeted, all-viral chemogenetic approach (as illustrated in Fig. 2A) (67). First, we injected a recombinant adeno-associated viral vector (AAV) for the CaMKII-driven expression of the recombinase Cre in the ventricle of the developing brain at the time of EBN and LBN neurogenesis, to achieve selective expression of Cre in isochronic neurons (Step 1 in Fig. 2A). Then, we performed a CA3-targeted local injection of an AAV carrying a construct for the Cre-dependent expression of Designer Receptors Exclusively Activated by Designer Drugs (DREADDs) in adult animals (*75*), to achieve selective expression of chemogenetic receptors in hippocampal EBNs and LBNs (Step 2 in Fig. 2A; Fig. 2B and (S4A). The all-viral approach was as efficient and selective as the BrdU approach in targeting hippocampal EBNs and LBNs (Fig. S4, B and C). Moreover, by targeting either the inhibitory (hM4Di) or excitatory (hM3Dq) DREADD to isochronic neurons, our approach was successful in preventing or inducing recruitment of EBNs and LBNs into c-Fos+ ensembles upon the systematic delivery of the DREADD ligand CNO, respectively (Fig. S4, D-I).

**Fig. 2.**
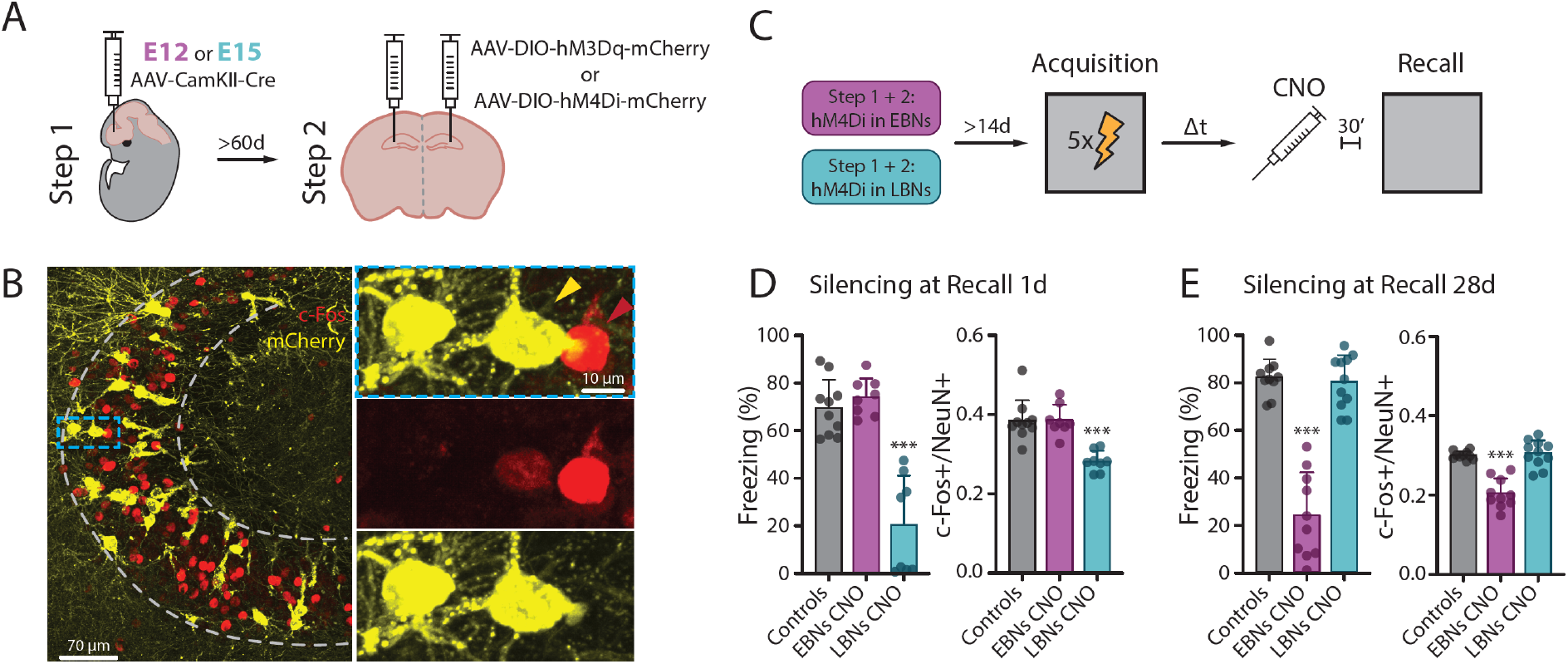
Timely recruitment of EBNs and LBNs at Recall is necessary for memory retrieval. **(A)** Schematic of the all-viral approach to target isochronic hippocampal neurons in the mouse brain. Step 1: Targeting Cre expression to isochronic neurons. Step 2: Targeting Cre-dependent expression of DREADDs (hM3Dq- and hM4Di-mCherry) to hippocampal EBNs and LBNs. Panels for Step 1 were created with BioRender.com. **(B)** c-Fos (red) and mCherry (yellow) expression in isochronic CA3 pyramidal neurons targeted by the all-viral method described in Fig. 2A. The inset (dashed blue box) is magnified on the right, where individual channels for c-Fos (red) and mCherry (yellow) are shown. The red arrow points to c-Fos+, mCherry-neurons, yellow arrow points to a c-Fos+, mCherry+ neuron. The scale bar is 70 µm for the overview image on the left, and 10 µm for the magnification on the right. Images are maximum intensity projections, 30X silicone objective. **(C)** Schematic of the experiment for the silencing of EBNs and LBNs at cFC Recall. Adult (>P60) mice where EBNs (magenta) or LBNs (cyan) had been targeted for the expression of the inhibitory version of the DREADD receptor (hM4Di), were subjected to the cFC protocol. 30 minutes before the Recall session, DREADD ligand CNO was injected *i.p.* to silence isochronic neurons. **(D)** Left: freezing during the exploration of the Training Context at Recall 1d in Control animals (N = 10, a mix of No Virus/No CNO; Step 1+ 2 Virus/No CNO; No Virus/CNO animals), and upon DREADD-mediated silencing of EBNs (N = 8) or LBNs (N = 8) via CNO injection. Each dot is a different mouse. Bars represent the mean, error bars the SD. Memory retrieval was significantly impaired when silencing was targeted to LBNs, but not when it was targeted to EBNs (one-way ANOVA, F(2, 23) = 35.82, p<0.0001. Dunnett’s multiple comparisons test to Controls: p>0.05 for EBNs silencing; p<0.0001 for LBNs silencing). Right: c-Fos expression in the CA3 network upon isochronic neurons silencing during Recall 1d. y axis: fraction of c-Fos+, NeuN+ neurons normalized by the total number of neurons in the CA3 network (NeuN+). Each dot is a different mouse. Bars represent the mean, error bars the SD. The size of the c-Fos+ ensemble recruited upon memory retrieval was significantly smaller than Controls when DREADDs were targeted to LBNs, but not when they were targeted to EBNs (one-way ANOVA, F(2, 23) = 17.44, p<0.0001. Dunnett’s multiple comparisons test to Controls: p>0.05 for EBNs silencing; p<0.0001 for LBNs silencing). **(E)** Left: freezing during the exploration of the Training Context at Recall 28d in Control animals (N = 10) and upon DREADD-mediated Silencing of EBNs (N = 10) or LBNs (N = 11) via CNO injection. Each dot is a different mouse. Bars represent the mean, error bars the SD. Memory retrieval was significantly impaired when silencing was targeted to EBNs, but not when it was targeted to LBNs (one-way ANOVA, F(2, 28) = 65.23, p<0.0001. Dunnett’s multiple comparisons test to Controls: p<0.0001 for EBNs silencing; p>0.05 for LBN silencing). Right: c-Fos expression in the CA3 network upon isochronic neurons silencing during Recall 28d. y axis: fraction of c-Fos+, NeuN+ neurons normalized by the total number of neurons in the CA3 network (NeuN+). Each dot is a different mouse. Bars represent the mean, error bars the SD. The size of the c-Fos+ ensemble recruited upon memory retrieval was significantly smaller than Controls when DREADDs were targeted to EBNs, but not when they were targeted to LBNs (one-way ANOVA, F(2, 28) = 37.65, p<0.0001. Dunnett’s multiple comparisons test to Controls: p<0.0001 for EBN silencing; p>0.05 for LBN silencing).

In a first set of experiments, we induced the expression of hM4Di in EBNs or LBNs to silence their activity and prevent their recruitment in memory ensembles at Recall (Fig. 2C). Silencing LBNs at Recall 1d significantly impaired memory retrieval, as indicated by the significantly lower levels of freezing displayed by the silenced group in comparison to controls (*i.e.* a pool of naïve and CNO-only injected animals, Fig. 2D, left), and reduced the size of the c-Fos+ ensemble established upon this experience in the CA3 network (Fig. 2D, right). Importantly, in animals in which a similar fraction of the total hippocampal network was silenced, but DREADD expression was restricted to EBNs instead of LBNs, levels of freezing and the size of the c-Fos+ ensemble at Recall 1d were comparable to controls (Fig. 2D). This indicated that the effects observed in the LBN-silenced group were not due to a general disruption of network activity but to the specific decrease in the activity of the Late Born subpopulation. Moreover, in LBN-but not in EBN-silenced mice, individual animal’s freezing levels were inversely correlated to the overall prevalence of DREADD-expressing neurons within the CA3 network (Fig. S5A), further suggesting a strong relationship between LBN silencing and memory retrieval impairment upon Recall 1d. In a mirrored fashion, silencing EBNs upon memory Recall 28d significantly impaired memory retrieval (Fig. 2E, left) and reduced the size of the CA3 c-Fos+ ensemble (Fig. 2E, right), while silencing LBNs at the same time did not produce significant effects. Hence, in EBN-but not LBN-targeted animals, freezing levels were inversely correlated to the overall prevalence of DREADD-expressing neurons within the CA3 network (Fig. S5B).

In a different set of experiments, we induced the expression of hM3Dq in EBNs or LBNs, and used chemogenetics to promote the recruitment of these subpopulations to c-Fos+ ensembles in an anachronistic fashion through the activation of EBNs at Recall 1d, and LBNs at Recall 28d (Fig. S6A). In both experiments, priming the recruitment of the “wrong” isochronic population impaired memory retrieval, as indicated by the significantly lower amount of freezing of the DREADD-expressing, CNO-injected animals in comparisons to controls (Fig. S6, B and C, left). Interestingly, such manipulations did not exert any effect on the size of the c-Fos+ ensemble activated upon recall (Fig. S6, B and C, right).

Taken together, targeted manipulation experiments revealed that the timely recruitment of EBNs or LBNs in c-Fos+ ensembles is crucial for memory retrieval at different stages of a memory’s lifetime, with each subpopulation’s activity being necessary at times when its recruitment into c-Fos+ memory ensembles is at its peak.

## LBN activity is necessary to support the long-term expression of EBN-dependent memory

Notably, silencing EBNs at times when LBN activity was necessary for retrieval (and vice versa) did not impair the expression of the fear memory, indicating that at Recall, changes in the activity in one isochronic population do not affect the other’s ability to support memory retrieval. Hence, we set out to determine the extent to which EBNs and LBNs might be mutually reliant on each other’s activity during the initial phases of memory formation.

First, we focused on mutual interactions during memory encoding. To reveal if activity from one isochronic subpopulation would affect the recruitment of the other into the c-Fos+ ensemble established at Acquisition, we used a double labelling approach through which, in the same animal, the expression of the inhibitory DREADD was targeted to one subpopulation while the other was labelled by the injection of BrdU (Fig. 3, A and B). Silencing either EBNs or LBNs at Acquisition successfully prevented the recruitment of DREADD-expressing neurons in the c-Fos+ ensemble established upon memory encoding, as expected (Fig. 3C). At the same time, levels of c-Fos expression in the BrdU-targeted subpopulation (LBNs and EBNs, respectively) were not different from controls (Fig. 3D). Moreover, on an animal-to-animal basis, there was no correlation between the extent of silencing in one isochronic subpopulation and the extent of recruitment of the other subpopulation (Fig. 3E). These data strongly indicate that the recruitment of EBNs and LBNs to c-Fos+ ensemble upon memory encoding occurs without mutual interactions between the two subpopulations, and that the establishment of the EBN- and LBN-dependent memory traces might be executed in parallel at Acquisition.

**Fig. 3.**
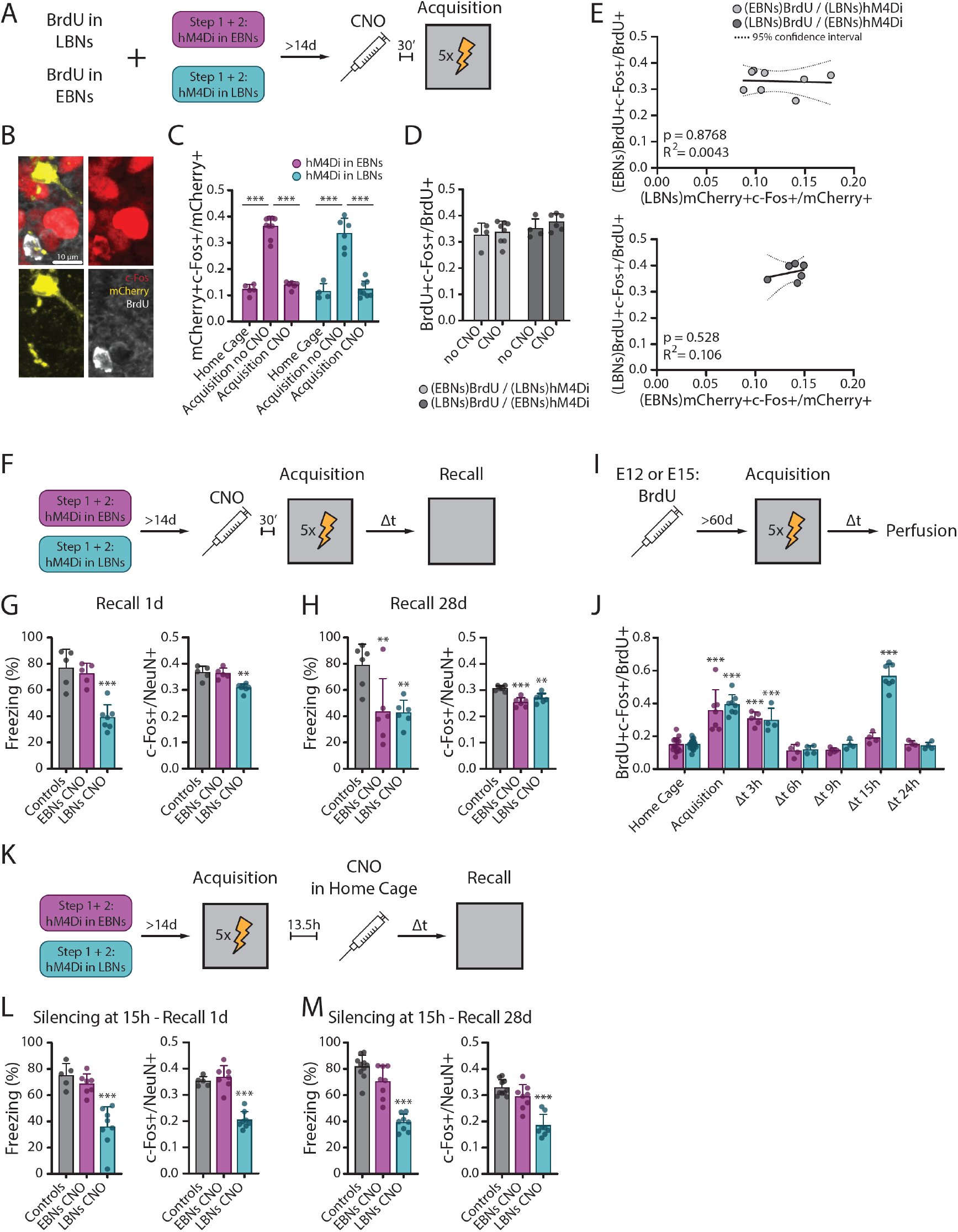
LBN activity during the early stages of memory processing is necessary for the expression of EBN-dependent memory. **(A)** Schematic for the combined approach targeting one isochronic subpopulation for the expression of the inhibitory DREADD receptor hM4Di, and the other subpopulation for the incorporation of BrdU. **(B)** c-Fos (red), mCherry (yellow), and BrdU (white) immunolabelling of CA3 neurons upon the combined approach described in Fig. 3A. Scale bar is 10 µm, images are maximum intensity projections, 30X silicone objective. **(C)** c-Fos expression in hM4Di-expressing EBNs (magenta) and LBNs (cyan) in Home Cage housed, hM4Di-expressing animals (EBNs: N = 5. LBNs: N = 4); hM4Di-expressing animals not injected with CNO upon cFC Acquisition (EBNs: N = 9. LBNs: N = 6); and hM4Di-expressing animals injected with CNO upon cFC Acquisition (EBNs: N = 6; LBNs: N = 8). y-axis: fraction of DREADD-expressing isochronic neurons (mCherry+) positive for c-Fos (mCherry+ c-Fos+/mCherry+). Each dot is a different mouse. Bars represent the mean, error bars the SD. CNO injection prevented the recruitment of the DREADD-targeted subpopulation in the c-Fos+ ensemble supporting memory encoding. For DREADD expression in EBNs: one-way ANOVA, F(2,17) = 57.34, p<0.0001. Tukey’s multiple comparisons test: Home Cage Vs no CNO, p<0.0001; Home Cage vs. CNO, p >0.05; no CNO vs. CNO, p<0.0001. For DREADD expression in LBNs: one-way ANOVA, F(2,15) = 31.65, p<0.0001. Tukey’s multiple comparisons test: Home Cage Vs no CNO, p<0.0001; Home Cage vs. CNO, p>0.05; no CNO vs. CNO, p<0.0001. **(D)** c-Fos expression in BrdU+ neurons of animals where EBNs were labelled with BrdU while LBNs expressed hM4Di (indicated as (EBNs)BrdU/(LBNs)hM4Di, light grey), or in animals where LBNs were labelled with BrdU while EBNs expressed hM4Di (indicated as (LBNs)BrdU/(EBNs)hM4Di, dark grey). Animals were either not injected ((EBNs)BrdU/(LBNs)hM4Di: N = 4. (LBNs)BrdU/(EBNs)hM4Di: N = 4) or injected with CNO ((EBNs)BrdU/(LBNs)hM4Di: N = 8. (LBNs)BrdU/(EBNs)hM4Di: N = 6) before cFC Acquisition. Y-axis: fraction of BrdU+ neurons expressing c-Fos (BrdU+ c-Fos+/BrdU+). Each dot is a different mouse. Bars represent the mean, error bars the SD. Silencing neurons belonging to one subpopulation of isochronic neurons did not prevent the recruitment of the non-targeted subpopulation in the c-Fos+ ensemble supporting memory encoding. For (EBNs)BrdU/(LBNs)hM4Di, CNO vs. no CNO: two-tailed student’s t-test, t(10) = 0.294, p>0.05. For (LBNs)BrdU/(EBNs)hM4Di, CNO vs. No CNO: two-tailed student’s t-test, t(8) = 1.182, p>0.05. **(E)** A simple linear regression was calculated to predict c-Fos expression in EBNs (top) or LBNs (bottom) based on the fraction of the CA3 network that expressed the DREADD receptor hM4Di when this was targeted to LBNs or EBNs, respectively, and CNO was injected at Acquisition. No significant regressions were found in either case. For (EBNs)BrdU / (LBNs)hM4Di upon CNO injection at Acquisition: F(1,6) = 0.026, p>0.05. For (LBNs)BrdU / (EBNs)hM4Di upon CNO injection at Acquisition: F(1,4) = 0.4756, p>0.05. Samples size as in Fig. 3D. Dotted lines express 95% confidence intervals. **(F)** Schematic for the silencing of EBNs and LBNs upon cFC Acquisition. **(G)** Left: freezing during the exploration of the Training Context at Recall 1d in Control animals (N = 5) and upon DREADD-mediated silencing of EBNs (N = 5) or LBNs (N = 7) at Acquisition. Each dot is a different mouse. Bars represent the mean, error bars the SD. Memory retrieval was significantly impaired when silencing was targeted to LBNs, but not when it was targeted to EBNs (one-way ANOVA, F(2, 14) = 21.77, p<0.0001. Dunnett’s multiple comparisons test to Controls: p>0.05 for EBNs silencing; p<0.0001 for LBNs silencing). Right: c-Fos expression in the CA3 network upon Recall 1d when isochronic neurons were silenced at Acquisition. Y-axis: fraction of c-Fos+, NeuN+ neurons normalized by the total number of neurons in the CA3 network (NeuN+). Each dot is a different mouse. Bars represent the mean, error bars the SD. The size of the c-Fos+ ensemble recruited upon memory retrieval was significantly smaller than Controls when DREADDs were targeted to LBNs, but not when they were targeted to EBNs (one-way ANOVA, F(2, 14) = 15.92, p = 0.0002. Dunnett’s multiple comparisons test to Controls: p>0.05 for EBNs silencing; p = 0.005 for LBNs silencing). **(H)** Left: freezing during the exploration of the Training Context at Recall 28d in Control animals (N = 6) and upon DREADD-mediated Silencing of EBNs (N = 6) or LBNs (N = 6) at Acquisition. Each dot is a different mouse. Bars represent the mean, error bars the SD. Memory retrieval was significantly impaired when silencing was targeted to either EBNs or LBNs (one-way ANOVA, F(2, 15) = 7.576, p = 0.0053. Dunnett’s multiple comparisons test to Controls: p = 0.0083 for EBNs silencing; p = 0.0075 for LBNs silencing). Right: c-Fos expression in the CA3 network upon Recall 28d when isochronic neurons were silenced at Acquisition. Y-axis: fraction of c-Fos+, NeuN+ neurons normalized by the total number of neurons in the CA3 network (NeuN+). Each dot is a different mouse. Bars represent the mean, error bars the SD. The size of the c-Fos+ ensemble recruited upon memory retrieval was significantly smaller than Controls when DREADDs were targeted to either EBNs or LBNs (one-way ANOVA, F(2, 15) = 17.42, p = 0.0001. Dunnett’s multiple comparisons test to Controls: p<0.0001 for EBNs silencing; p = 0.002 for LBNs silencing). **(I)** Schematic of the cFC consolidation experiment on BrdU pulse-chased animals. During the Acquisition session, adult (>P60) mice who had been pulse-chased during development through an injection of BrdU at either E12 (EBNs labelled, magenta) or E15 (LBNs labelled, cyan) were introduced into the conditioning chamber. cFC protocol was then performed as in Fig. 1A. After a variable delay from Acquisition (Δt, from 3 to 24 hours), mice were taken from their Home Cage and tissue was processed for further analysis. **(J)** Time course of expression of c-Fos in CA3 EBNs (magenta) and LBNs (cyan) in the hours following the encoding of a cFC memory. Each dot is an animal. Bars represent the mean, error bars the SD. c-Fos expression in EBNs exhibited a peak that started at Acquisition and remained elevated for 3 hours after, to go back to baseline by 6h and up to 24h thereafter (one-way ANOVA, F(6, 38) = 15.45, p<0.0001. Dunnett’s multiple comparisons test to Home Cage EBNs: p<0.0001 for Acquisition and Δt 3h; p>0.05 for Δt=6h, 9h, 15h, and 24h). LBNs instead exhibited two distinct waves of c-Fos expression, one with a peak sustained between Acquisition and 3h, and a second centred around 15h after memory encoding (one-way ANOVA, F(6, 45) = 97.29, p<0.0001. Dunnett’s multiple comparisons test to Home Cage LBNs: p<0.0001 for Acquisition, Δt 3h, and Δt 15h; p>0.05 for Δt = 6h, 9h, and 24h). At least 4 animals per group, no repeated measures. **(K)** Schematic for the silencing of EBNs and LBNs upon the consolidation of a cFC memory. Adult (>P60) mice where EBNs (magenta) or LBNs (cyan) had been targeted with the all-viral method described in Fig. 2A for the expression of the inhibitory version of the DREADD receptor, were subjected to the cFC protocol as described in Fig. 1A. 13.5 hours after the Acquisition session, and while animals were housed in their home cage, DREADD ligand CNO was injected *i.p.* to silence isochronic neurons. **(L)** Left: freezing during the exploration of the Training Context at Recall 1d in Control animals (N = 5) and upon DREADD-mediated silencing of EBNs (N = 7) or LBNs (N = 8) during the second wave of c-Fos expression after Acquisition (*i.e.* 15 hours post-Acquisition, CNO injection at 13.5h). Each dot is an animal. Bars represent the mean, error bars the SD. Memory retrieval was significantly impaired when silencing was targeted to LBNs, but not when it was targeted to EBNs (one-way ANOVA, F(2, 17) = 21.59, p<0.0001. Dunnett’s multiple comparisons test to Controls: p>0.05 for EBNs silencing; p<0.0001 for LBNs silencing). Right: c-Fos expression in the CA3 network upon Recall 1d when isochronic neurons were silenced during the second wave of c-Fos expression after Acquisition. Y-axis: fraction of c-Fos+, NeuN+ neurons normalized by the total number of neurons in the CA3 network (NeuN+). Each dot is a different mouse. Bars represent the mean, error bars the SD. The size of the c-Fos+ ensemble recruited upon memory retrieval was significantly smaller than Controls when DREADDs were targeted to LBNs, but not to EBNs (one-way ANOVA, F(2, 17) = 47.82, p<0.0001. Dunnett’s multiple comparisons test to Controls: p>0.05 for EBNs silencing; p<0.0001 for LBNs silencing). **(M)** Left: freezing during the exploration of the Training Context at Recall 28d in Control animals (N = 10) and upon DREADD-mediated silencing of EBNs (N = 8) or LBNs (N = 8) during the second wave of c-Fos expression after Acquisition (*i.e.* 15 hours post-Acquisition, CNO injection at 13.5h). Each dot is an animal. Bars represent the mean, error bars the SD. Memory retrieval was significantly impaired when silencing was targeted to LBNs, but not when it was targeted to EBNs (one-way ANOVA, F(2, 23) = 46.30, p<0.0001. Dunnett’s multiple comparisons test to Controls: p>0.05 for EBNs silencing; p<0.0001 for LBNs silencing). Right: c-Fos expression in the CA3 network upon Recall 28d when isochronic neurons were silenced during the second wave of c-Fos expression after Acquisition. Y-axis: fraction of c-Fos+, NeuN+ neurons normalized by the total number of neurons in the CA3 network (NeuN+). Each dot is a different mouse. Bars represent the mean, error bars the SD. The size of the c-Fos+ ensemble recruited upon memory retrieval was significantly smaller than Controls when DREADDs were targeted to LBNs, but not to EBNs (one-way ANOVA, F(2, 23) = 30.58, p<0.0001. Dunnett’s multiple comparisons test to Controls: p>0.05 for EBNs silencing; p<0.0001 for LBNs silencing).

If EBNs and LBNs were to give rise to independent memory traces, then a failure in encoding one trace should impact memory retrieval only during times when that specific trace is utilized, but not when the other subpopulation supports recall. To test this, we evaluated how silencing isochronic neurons during Acquisition affected memory retrieval at Recall (Fig 3F). Silencing LBNs (but not EBNs) during Acquisition impaired memory retrieval at Recall 1d. This impairment was evident through significantly reduced freezing levels in LBN-silenced animals compared to controls (Fig. 3G, left), a smaller size of the CA3 c-Fos+ ensemble in these subjects (Fig. 3G, right), and an inverse correlation between the prevalence of DREADD-expressing neurons within the CA3 network and freezing levels specifically in LBN-silenced animals (Fig. S7A). However, a different outcome could be observed when either isochronic subpopulation was silenced during Acquisition, and memory retrieval was assessed 28 days later. Then, the expression of fear memory was notably hampered (Fig. 3H, left), with the CA3 c-Fos+ ensemble at Recall being smaller than controls (Fig. 3H, right), regardless of whether EBNs or LBNs were silenced during Acquisition. Thus, while both EBN and LBN activity during Acquisition was necessary for the expression of the fear memory at times when each subpopulation’s recruitment was maximal, the successful expression of the EBN-dependent memory hinged on LBNs’ activation during memory encoding. Yet, freezing levels at the 28-day Recall inversely correlated with DREADD expression in EBN-targeted (but not LBN-targeted) animals (Fig. S7B). This indicates that EBN activity maintained a closer connection to memory retrieval during remote recalls.

In a second set of experiments, we focused on mutual interactions during the initial phases of memory consolidation. For this, we first systematically monitored the expression of c-Fos among EBNs and LBNs in mice that were not re-exposed to the training context during the hours after memory encoding (*i.e.* in mice that were housed in their home cage at any time after Acquisition, Fig. 3I). Here too, EBNs and LBNs exhibited distinct dynamics of c-Fos expression. While c-Fos expression in EBNs was elevated for a few hours following Acquisition, and comparable to Home Cage controls by 6 – and up to 24 – hours after (Fig. 3J), LBNs exhibited two distinct waves of increased c-Fos expression. The first recapitulated EBNs, with a peak after encoding that was back to baseline by 6 hours. The second wave instead peaked around 15 hours and was back to baseline by 24 (Fig. 3J), consistent with previous observations that identified a protein synthesis- and dopamine signalling-dependent second window of consolidation at comparable times (Fig. S8A) (*76, 77*). Strikingly, LBN, but not EBN, activity during this window was necessary for the establishment of c-Fos+ ensembles and correct retrieval of the cFC memory at Recall 1d and 28d after Acquisition (Fig. 3, K-M), when freezing levels were inversely correlated to the expression of DREADDs in LBN-, but not EBN-targeted animals (Fig. S8, B and C). As a control, silencing both EBNs or LBN at times when neither subpopulations nor the whole hippocampal network exhibited a significant difference in c-Fos expression in comparison to controls (*i.e.* 9 hours after memory encoding, Fig. 3J and (S8A) exerted no effect on memory retrieval or c-Fos expression upon Recall (Fig. S8, D and E).

Taken together, our results indicate that the recruitment of both Early- and Late-Born neurons to c-Fos+ ensembles at Acquisition operates independently, with no mutual interactions between the subpopulations. Moreover, EBN and LBN activity at Acquisition is necessary for the retrieval of the fear memory when each subpopulation exhibits peak c-Fos expression at Recall. Finally, LBN activity at Acquisition or during a specific window of memory consolidation is critical for the long-term expression of the EBN-dependent memory.

## LBNs-to-EBNs recruitment shift defines the temporal boundaries for a transient window of enhanced plasticity

Lastly, we explored the functional implications of the shift in recruitment from Late- to Early-Born neurons for hippocampus-dependent memory processes. Previous work revealed that in the hippocampus, distinct cohorts of isochronic neurons are endowed with distinct activity dynamics when it comes to ensemble firing, intrinsic properties, locking to oscillations, and tuning to space (*58, 71*). Moreover, neurons with late-born firing dynamics take part in neuronal ensembles that, upon learning, exhibit a higher degree of plasticity than neurons with early-born firing patterns (*62, 68, 78*). Strikingly, with time a memory exhibits a different potential for plasticity, being more prone to be updated and having heightened susceptibility to pharmacological manipulations shortly after its encoding than at remote times (*33, 36*). Thus, we hypothesized that the differential recruitment of LBNs and EBNs in memory ensembles might underpin a memory’s potential for plasticity. For this, we explored two learning paradigms where an animal’s behaviour was a function of its ability to integrate information across multiple learning episodes.

In a first line of experiments, we investigated mechanisms of memory plasticity associated with the ability to reinforce learned associations. For this, we leveraged the finding that in a cFC protocol when all USs are experienced within the same session, like the one implemented so far in the study (*i.e.* a “Mass training” regime), the freezing behaviour of mice during Recall directly relates to how many USs they experience at Acquisition (*79*). Hence, when mice were presented with a train of 2-US at Acquisition, the amount of freezing expressed at Recall was halved compared to the presentation of a train of 5-US (Fig. S9, A and B). We then reasoned that if mice were able to integrate information across multiple learning episodes, in a protocol where they were exposed to two separate 2-US Acquisition sessions, the re-exposure to the US during the second session should reinforce the CS-US association learned in the first session, as supported by research on spaced training models (*80, 81*). This could mean that mice initially showing low freezing after the first 2-US Acquisition session, might exhibit significantly higher levels of freezing after the second 2-US Acquisition session.

Thus, in the Spaced training cFC task (Fig. 4A), animals that underwent a 2-US Acquisition session (“Training” session), experienced a second 2-US Acquisition session (“Re-Training” Session) in the same context and at variable temporal delays from the first (“Inter-Session Interval”, ISI). Their memory was tested both during the initial part of the Re-Training session (*i.e.* before the presentation of the US, to assess the formation of a weak-fear memory after Training) and during a Recall session performed 5 days from Training (to assess changes in memory strength after Re-Training). Mice subjected to spaced training with an ISI of 2 days or shorter could successfully strengthen their weak-fear memory into a strong-fear memory throughout the experiment, as indicated by a pronounced increase in freezing over sessions up to Recall (Fig 4B). However, for an ISI of 3 days or longer, mice failed to do so, and expressed similar levels of freezing across sessions (Fig. 4B). Such differences were stable at 28 days after encoding (Fig. S9C). Thus, we concluded that cFC memories encoded in a spaced training paradigm can be updated for the reinforcement of a learned association between a CS and the US during a temporally restricted plasticity window lasting up to the second day from the initial learning episode.

**Fig. 4.**
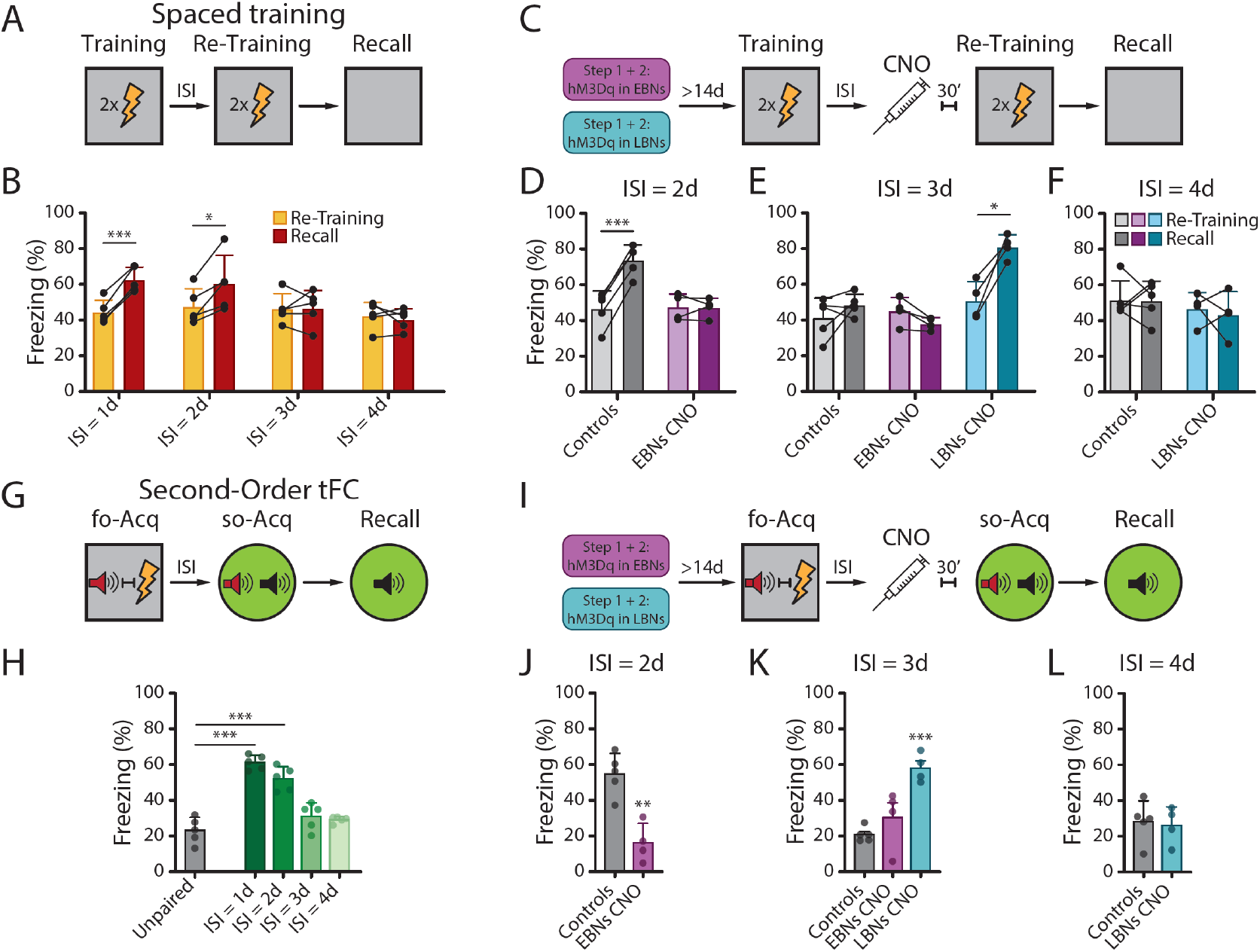
Integration of learning episodes into cohesive memories is modulated by EBNs and LBNs recruitment to c-Fos+ ensembles. **(A)** Schematic of the spaced cFC protocol. During the Training session, adult (>P60) mice were introduced into the conditioning chamber, and let explore the Training Context for 4 minutes before receiving a train of 2 shocks at 30-second intervals, before being brought back to their home cage. A second training session (“Re-Training”) was performed with a delay of a variable number of days from Training (ISI, going from 1 to 4 days). Distinct cohorts of animals were trained in the task for ISI = 1d, 2d, 3d, and 4d. For all cohorts, the Recall session was performed on day 5 from Training, during which mice were reintroduced back into the Training Context to assess memory retrieval. **(B)** Freezing for animals exploring the Training context during the Re-Training (yellow, % of time spent freezing before the delivery of the shocks) and Recall (red) sessions of the spaced cFC task. For each ISI group, each dot is a different mouse and each mouse is tested longitudinally across Re-Training and Recall sessions (connecting lines). Bars represent the mean, error bars the SD. Animals who trained in the spaced cFC protocol (N = 5 animals for each group) could combine the two learning episodes for the reinforcement of the CS/US association if the Re-Training Session was performed with an ISI≤2d day from Training, as indicated by the significantly elevated levels of freezing at Recall in comparison with those at Re-Training (for ISI = 1: paired t-test, t(4) = 10.61, p = 0.0004. For ISI = 2: paired t-test, t(4)=3.820, p = 0.0188). For ISI≥3d days, freezing levels expressed after the first and second learning episodes were comparable, suggesting that the Re-Training session did not lead to a reinforcement of the learned CS/US association (for ISI = 3: paired t-test, t(4) = 0.071, p>0.05. For ISI = 4: paired t-test, t(4)=0.662, p>0.05). **(C)** Schematic of the experiment for the activation of isochronic neurons upon the spaced cFC protocol. Adult (>P60) mice where EBNs (magenta) or LBNs (cyan) had been targeted with the all-viral method described in Fig. 2A for the expression of the excitatory version of the DREADD receptor hM3Dq, were subjected to the spaced cFC protocol as described in Fig. 4A. 30 minutes before the Re-Training session, DREADD ligand CNO was injected *i.p.* to activate isochronic neurons. **(D)** Percentage of time spent freezing during the exploration of the Training context upon the Re-Training (light shading, % of time spent freezing before the delivery of the shocks) and Recall (dark shading) sessions of the spaced cFC task in Control mice and upon the activation of EBNs during Re-Training, which was performed with ISI = 2d from Training. Each dot is a different mouse and each mouse is tested longitudinally across Re-Training and Recall sessions (connecting lines). Bars represent the mean, error bars the SD. In control animals, the spaced cFC task with ISI = 2d led to the reinforcement of the learned CS/US association, consistent with data presented in Fig. 4B (N = 5 animals. Paired t-test, t(4) = 20.89, p<0.0001). Activation of EBNs during Re-Training (N = 4) impaired the integration of information between the two training sessions, with a consequent expression of a weak-fear memory at Recall (paired t-test, t(3) = 0.164, p>0.05). **(E)** Percentage of time spent freezing during the exploration of the Training context upon the Re-Training (light shading, % of time spent freezing before the delivery of the shocks) and Recall (dark shading) sessions of the spaced cFC task in Control mice and upon the activation of EBNs or LBNs during Re-Training, which was performed with ISI = 3d from Training. Each dot is a different mouse and each mouse is tested longitudinally across Re-Training and Recall sessions (connecting lines). Bars represent the mean, error bars the SD. In controls, the spaced cFC task with ISI = 3d did not lead to the reinforcement of the learned CS/US association, consistent with data presented in Fig. 4B (N = 5, grey, paired t-test, t(4) = 1.249, p>0.05). Activation of EBNs during re-training (N = 4, magenta) did not produce any significant effect on the ability to integrate information across the two training sessions, as indicated by similar levels of freezing at Re-Training and Recall (paired t-test, t(3) = 1.205, p>0.05). In stark contrast, the activation of LBNs during Re-Training (N = 4, cyan) led to successful integration for the reinforcement of the learned association and significantly higher freezing at Recall (paired t-test, t(3) = 5.404, p=0.0124). **(F)** Percentage of time spent freezing during the exploration of the Training context upon the Re-Training (light shading, % of time spent freezing before the delivery of the shocks) and Recall (dark shading) sessions of the spaced cFC task in Control mice and upon the activation of EBNs during Re-Training, which was performed with ISI = 4d from Training. Each dot is a different mouse and each mouse is tested longitudinally across Re-Training and Recall sessions (connecting lines). Bars represent the mean, error bars the SD. In control animals, the spaced cFC task with ISI = 4d did not lead to the reinforcement of the learned CS/US association, consistent with data presented in Fig. 4B (N = 5 animals. Paired t-test, t(4) = 0.184, p>0.05). Similarly, in animals where LBNs were activated during Re-Training, this second learning episode failed to reinforce the learned association (N = 4, paired t-test, t(3) = 0.461, p>0.05). **(G)** Schematic of the second order tFC protocol. During the first-order Acquisition session (fo-Acq), adult (>P60) mice were introduced into the conditioning chamber, let explore the Training Context for 3 minutes before receiving a train of 6 Tone-CS/Trace/Shock pairing presentations (as in the tFC paradigm, see Methods), before being brought back to their home cage. A second-order Acquisition session (so-Acq) was performed at a delay of a variable number of days from the fo-Acq (ISI, going from 1 to 4 days). Distinct cohorts of animals were trained in the task for ISI = 1d, 2d, 3d, and 4d. During the so-Acq session, mice were introduced into a novel context (Context B) and let explore for 3 minutes before being exposed to a series of 5 pairings between the second-order CS (so-CS, white noise, black symbol in the schema) and the first-order CS (fo-CS, red symbol in the schema, Tone used during fo-Acq). In the Unpaired Controls group, no fo-CS was presented during the so-Acq session, to assess baseline levels of freezing to the White Noise in the absence of the formation of an inference. For all cohorts, the Recall session was performed on day five from fo-Acq, during which mice were reintroduced in Context B and freezing was quantified upon the presentation of the so-CS. **(H)** Freezing during the Recall session of the so-tFC task. Each dot is a different mouse. Bars represent the mean, error bars the SD. N = 5 animals for each group. Animals where the so-Acq session was performed at ISI≤2d from the fo-Acq session could successfully form an inference between the so-CS and the fo-CS, as indicated by the significantly higher levels of freezing to the presentation of the so-CS during Recall in comparison to the Unpaired Controls. In contrast, animals where the so-Acq was performed at ISI≥3 days exhibited levels of freezing that were comparable to the Unpaired Controls. One-way ANOVA, F(4, 20) = 38.18, p<0.0001. Tukey’s multiple comparisons test: Unpaired Controls vs. ISI = 1d, p<0.0001; Unpaired Controls vs. ISI = 2d, p<0.0001; ISI = 1d vs. ISI = 3d, p<0.0001; ISI = 2d vs. ISI = 3d, p = 0.0002; ISI = 2d vs. ISI=4d, p<0.0001; p>0.05 for all other comparisons). **(I)** Schematic of the experiment for the activation of isochronic neurons upon the so-tFC protocol. Adult (>P60) mice where EBNs (magenta) or LBNs (cyan) had been targeted with the all-viral method described in Fig. 2A for the expression of the excitatory version of the DREADD receptor hM3Dq, were subjected to the Second-Order tFC protocol as described in Fig. 4G. 30 minutes before the so-Acq session, DREADD ligand CNO was injected *i.p.* to activate isochronic neurons. **(J)** Freezing during the Recall session of the so-tFC task upon activation of EBNs during so-Acq performed at ISI = 2d. Each dot is a different mouse. Bars represent the mean, error bars the SD. While in Controls (N = 5) the so-tFC task with ISI = 2d led to the formation of a second-order association between the so-CS and fo-CS as described in Fig. 4H, the activation of EBNs during so-Acq (N = 4) session impaired such process (unpaired t-test, t(7) = 5.057, p = 0.0015). **(K)** Freezing during the Recall session of the so-tFC task upon activation of isochronic neurons during so-Acq performed at ISI = 3d. Each dot is a different mouse. Bars represent the mean, error bars the SD. While in both Controls (N = 5, grey) and upon the activation of EBNs during the so-Acq session (N = 4, magenta), the so-tFC task with ISI = 3d did not lead to the formation of an inference, activation of LBNs during so-Acq (N = 4, cyan) led to the formation of a second-order association between so-CS and fo-CS (one-way ANOVA, F(2, 10) = 14.78, p = 0.001. Dunnett’s multiple comparison test to Controls: p>0.05 for EBNs activation, p = 0.0006 for LBNs activation). **(L)** Freezing during the Recall session of the so-tFC task upon activation of LBNs during so-Acq performed at ISI = 4d. Each dot is a different mouse. Bars represent the mean, error bars the SD. Neither controls (N = 5) nor LBNs-Activated animals (N = 4) were able to integrate information across training sessions for the formation of a second-order association between so-CS and fo-CS (unpaired t-test, t(7) = 0.30, p>0.05).

Strikingly, the time course of this phenomenon recapitulated the shift in the recruitment of Late-versus Early-Born neurons into c-Fos+ ensembles after Acquisition (Fig. 1, D and E), as well as the reactivation profile of Late- and Early-Born Engram Neurons (Fig. 1J). Thus, we tested whether the differential recruitment of EBNs and LBNs upon re-exposure to the training context in the Re-Training session might underpin the ability to integrate information across learning episodes. For this, we used the all-viral approach to induce the expression of hM3Dq in EBNs and LBNs, and induced the activation of these neurons by CNO injection upon Re-Training (Fig. 4C). In the first set of experiments, we activated EBNs when Re-Training was performed with ISI = 2d (akin to Recall d2 in Fig. 1, D and J), *i.e.* at times when c-Fos expression in EBNs is low, the reactivation of EB Engram neurons is at chance levels, and the initial weak-fear memory encoded upon Training is in its plasticity window. EBN-stimulated mice failed to use the Re-Training session to reinforce the association between CS and US, as demonstrated by levels of freezing at Recall that were in line with those expressed after Training and lower than controls (Fig. 4D). This result suggested that biasing the c-Fos+ ensemble towards EBNs during a memory’s plasticity window hampers the memory’s ability to be reinforced upon a second learning episode. In a second set of experiments, we activated either EBNs or LBNs when Re-Training was performed with ISI = 3d (akin to Recall d3 in Fig. Fig. 1, D and J), *i.e.* at times when c-Fos expression in EBNs is high but at baseline levels in LBNs, both EB and LB Engram neurons are significantly reactivated, but the weak-fear memory encoded at Training is barely past its plasticity window. In these experiments, while EBN-stimulated mice, just like controls, could not reinforce the initial CS-US association upon Re-Training, LBN-stimulated animals exhibited levels of freezing at Recall that were significantly higher than those exhibited after Training (Fig. 4E). Thus, priming the c-Fos+ ensemble towards enrichment in LBNs, but not EBNs, upon Re-Training performed just after the end of a memory’s plasticity window boosts the animal’s ability to reinforce a learned association through a second learning episode. Last, we investigated whether the stimulation of LBNs during Re-Training could lead to successful reinforcement at any time past the plasticity window. To this end, we chemogenetically activated LBNs when Re-Training was performed with ISI = 4d (akin to Recall d4 in Fig. 1, D and J), *i.e.* at times when c-Fos expression in LBNs is low, the reactivation of LB Engram neurons is at chance levels, and the initial weak-fear memory encoded at Training is days past its plasticity window. LBN-stimulated mice failed to use the Re-Training session to reinforce the association between CS and US, as demonstrated by levels of freezing at Recall that were similar to those expressed after Training and in line with Controls (Fig. 4F). This result suggested that the possibility to boost memory plasticity by biasing the c-Fos+ ensemble towards LBNs is temporally restricted to a short period after memory encoding.

In a different line of experiments, we investigated mechanisms of memory plasticity associated with the ability to use previously acquired information to make inferences (82–84). For this, we trained animals in a second-order Fear Conditioning paradigm (Fig. 4G). In this task, mice first learn to associate a first-order CS (fo-CS) with a US (a foot shock) during the first-order Acquisition session (fo-Acq). Then, during a second-order Acquisition session (so-Acq), the fo-CS is repeatedly presented together with a second, neutral conditioning stimulus (second-order CS, so-CS). Upon repeated presentation of the fo-CS / so-CS pairing, the presentation of so-CS in isolation during a Recall session elicits a significant fear response even though so-CS and US have never been experienced in the same session, revealing the encoding of an inferential association (Fig. 4G). In our experiment, we used the tFC protocol to create a hippocampus-dependent association between a tone (fo-CS) and the foot shock (US), and paired the tone with white noise (so-CS) during a so-Acq session performed at a variable ISI from the fo-Acq. Here, too, we hypothesized that the ability to make inferences might be a function of the temporal delay between training sessions, similar to what we observed in the spaced cFC task. Mice that underwent a so-cFC paradigm where fo-Acq and so-Acq sessions were performed with ISI≤2d froze significantly more to the presentation of the so-CS at Recall than animals in which the so-CS was never paired to the fo-CS (“Unpaired Controls”, animals that were exposed only to the so-CS during the so-Acq session, see Methods) (Fig. 4H). In stark contrast, mice did not learn the second-order association for ISI≥3d (Fig. 4H). These data therefore suggested that, as for the ability to reinforce a previously established CS-US association upon the presentation of a second learning episode, the ability to make inferences was a function of the temporal delay between Acquisition sessions. This prompted us to investigate if such ability could be modulated by the recruitment of Late-versus Early-Born isochronic neurons during the second-order Acquisition session. Hence, we followed the same logic of previous experiments, and chemogenetically primed EBNs and LBNs for their recruitment into the c-Fos+ ensembles during the so-Acq session when this was performed with a delay of 2, 3, or 4 days from the fo-Acq (Fig. 4I). In line with the spaced cFC experiments, neither the activation of EBNs nor the activation of LBNs upon the Second-Order session impaired the recall of the First-Order association memory, independently on the day when such association was tested (Fig. S9, D and E). However, the effects of the forced recruitment of EBNs or LBNs on the ability to create an inference depended on the ISI at which these two conditioning sessions were performed. Biasing the c-Fos+ ensemble towards EBNs during the so-Acq session for ISI = 2d prevented the formation of an inferential association, as indicated by the significantly lower freezing of CNO-injected, EBN-targeted animals vs. Controls at Recall (Fig. 4J and Fig. S9F, left). A similar manipulation on EBNs performed during the so-Acq session for ISI = 3d did not produce any significant effect compared to controls, which already did not show significant freezing to the presentation of so-CS at Recall (Fig. 4K and Fig. S9F, centre). Conversely, the activation of LBNs during the so-Acq session for ISI = 3d favoured the formation of an inferential association in conditions where this association would normally not be established, as indicated by the significantly higher freezing of CNO-injected, LBN-targeted animals vs. Controls during Recall (Fig. 4K and Fig. S9F, centre). Last, the activation of LBNs during the so-Acq session for ISI = 4d did not produce any significant effect compared to Controls, which already did not show freezing to the presentation of the so-CS at Recall (Fig. 4L and Fig. S9F, right).

Taken together, these results suggest that for a short window of time after a memory’s encoding, mice can exploit a second learning episode for the reinforcement of previously learned associations or the establishment of inferential associations. Moreover, manipulating the recruitment of LBNs and EBNs in c-Fos+ ensembles established during the second learning episode has the potential to modulate the temporal boundaries of such plasticity window for the integration of information across multiple learning episodes.

## Discussion

Our experiments reveal that the encoding of a fear-association memory results in the simultaneous and parallel emergence of two distinct memory traces within the mouse hippocampal network. These traces are instantiated in developmentally-defined subpopulations of early- and late-born principal neurons, and follow distinct reactivation trajectories post-encoding. In support of this, we show that late-born neurons are preferentially recruited to memory ensembles when a memory is recalled during a short window of time after encoding, during which LBN activity is necessary for memory retrieval. Such enrichment progressively shifts towards stable and sustained preferential recruitment of early-born neurons, whose activity is necessary for retrieval at longer delays. Thus, we discover a novel mechanism underlying a progressive reorganization of memory ensembles supporting the encoding, consolidation, and retrieval of fear-association memory over time, whose logic is grounded in the alternative recruitment of developmentally-defined subpopulations of principal neurons. We further demonstrate that memory expression at remote times depends on LBN activity at acquisition and during a specific window of consolidation. This evidence reveals that a dynamic interplay between isochronic subpopulations is crucial for a memory’s evolution and persistence over time. Last, we show that the systematic shift in recruitment from late- to early-born neurons to memory ensembles happens concomitantly to a previously unidentified transient period after memory encoding, when animals can combine information from multiple learning episodes to reinforce previously learned associations or make inferences. Through manipulation experiments, we demonstrate that modulating the degree of recruitment of early- or late-born neurons during a second learning episode was sufficient to modulate the animal’s performance in reinforcement and second-order learning tasks. Our findings therefore indicate that ensemble dynamics for the divergent recruitment of early- and late-born neurons have the potential to modulate the timing and extent of a transient plasticity window when the integration of information across multiple learning episodes into cohesive memories is facilitated.

While the time- and experience-dependent reorganization of neuronal ensembles supporting specific experiences, a phenomenon known as representational drift, has challenged prevailing theories of memory persistence, we show that for hippocampus-dependent fear association memories, ensemble reorganization is not disruptive of memory processes. Rather, the recruitment of distinct subpopulations of neurogenesis-defined neurons in a timely manner is necessary for the recall of hippocampal-dependent memories. Notably, silencing EBNs or LBNs specifically at times matching their peak recruitment in c-Fos+ ensemble, or artificially activating these subpopulations outside their normal recruitment periods, impaired memory retrieval. A seed for ensemble reorganization is already established during Acquisition through the parallel emergence of distinct developmentally-defined memory traces, and is driven by their alternative reactivation at recall, raising the question of how this differential recruitment might be achieved. One possible biological mechanism underlying EBNs and LBNs’ divergent trajectories could lie in the existence of different presynaptic networks, where the activation of distinct brain areas upon recent and remote recall might create a bias towards the recruitment of specific birth-dated neurons in the hippocampus (*28, 41, 44, 45*). Alternatively, the two isochronic subpopulations might be endowed with distinct plasticity properties which define unique trajectories for the emergence, reactivation, and permanence of the memory traces they support (*58*). Thus, we show that ensemble dynamics grounded in the alternative reactivation of developmentally-defined, parallel-emerging memory traces underpin the persistence of hippocampus-dependent fear memories over time. Although focusing on the recruitment dynamics of isochronic subpopulations that are born at the two extremes (*i.e.* start and end) of the CA3 neurogenesis period allowed us to discover parallel memory traces associated with these subpopulations, whether these traces are two of many, and similar or graded dynamics apply to neurons born at intermediate times during neurogenesis, is left for further investigation.

An important question is also whether and how EBNs and LBNs interact with each other to support memory dynamics. Our data indicate that the two subpopulations operate independently when allocating neurons to a c-Fos+ ensemble established upon memory encoding. Nevertheless, memory expression at remote times hinges on the activity of LBNs both during acquisition and within a specific consolidation window. Thus, offline reactivation of the transient memory trace supported by the activity of late-born neurons plays an instructive role in the maintenance and expression of the EBN-dependent memory. Such instructive function is unidirectional, given that silencing EBNs did not affect, at any time, the ability of LBNs to support memory retrieval, and might be underpinned by either local connectivity motifs or network interactions.

At the level of the local network, while isochronic principal neurons connect preferentially with each other across hippocampal subdivisions (*66, 85*), local computations via structured connectivity on interneurons might explain how LBN activity influences the long-term expression of the EBN-dependent memory trace. Previous reports have highlighted a connectivity motif where putative late-born pyramidal neurons exert a stronger excitatory drive on Parvalbumin-expressing (PV+) interneurons compared to putative EBNs, which receive greater inhibition from these interneurons (*60, 86*). Strikingly, the time course of LBN recruitment aligns with the dynamics of hippocampal PV+ network plasticity, which is known to be critical for the consolidation and long-term expression of fear memories (*77, 86, 87*). Therefore, a tight link between LBNs and PV+ network plasticity might underpin the instructive role played by late-born neurons on memory dynamics.

Alternatively, network-based computations might create an indirect route by which LBNs might influence EBN function in memory processing. Such a route might impinge on non-reciprocal communication between hippocampal LBNs, prefrontal ensembles, and hippocampal EBNs, and on the distinct propensity of the two subpopulations to be reactivated during offline events associated with memory consolidation (*70, 88, 89*). Within this context, network-based interactions between a “fast” learning system based on LBNs, and a “slow” system based on EBNs, where the first rapidly encodes a transient memory trace that drives the slow maturation of a permanent one, aligns with predictions formulated by the Complementary Learning Systems (CLS) theory (*90–92*). Accordingly, the fast system might quickly encode representations for unique experiences, while the slow system might gradually identify overarching patterns to form generalized knowledge structures (schemas). Existing evidence points to the hippocampus’s involvement in both computations, supporting both the rapid encoding of individual episodes and the construction of predictive models from various experiences (*93, 94*). The distinct roles of EBNs and LBNs in these processes remain to be determined.

Lastly, our experiments reveal that the ability to integrate information between related experiences for the reinforcement of learned associations or the creation of inferences decays as a function of the time elapsed between learning episodes. Strikingly, such decay recapitulates the temporal dynamics characterizing the shift from the preferential recruitment of LBNs to EBNs in memory ensembles. This observation led us to hypothesize that learning might define the opening of a temporally restricted plasticity window when multiple related experiences can be integrated into cohesive memories. In this framework, ensemble reorganization underpins the closure of such a window. In support of this hypothesis, we demonstrate that prematurely promoting the recruitment of EBNs into memory ensembles, during times when mice are physiologically capable of integrating information across experiences, disrupts this process. In stark contrast, enhancing the recruitment of LBNs during periods when mice typically cannot integrate across episodes improves their ability to do so, but this enhancement is only effective if it occurs within a few days after the initial learning episode. Therefore, understanding the cellular mechanisms of memory uncovered here holds the promise of enabling the deliberate reprogramming of memory dynamics. Notably, a failure in boosting memory plasticity upon LBN stimulation came at times when Late-Born Engram neuron reactivation was at chance levels, suggesting that stimulating LBNs to promote memory plasticity is most effective when the LBN-dependent memory trace can still be reactivated. In this framework, the transient LBN-dependent trace may function as a memory buffer for recently acquired information, facilitating the coherent processing of information acquired over multiple learning episodes into memories, and their subsequent routing to different memory-processing networks for long-term persistence (*4*).

In conclusion, our experiments demonstrate that early- and late-born hippocampal neurons exert distinct temporal and functional contributions to the processing and dynamics of fear-association memories. LBNs facilitate the establishment and reactivation of a transient memory trace upon learning, which might underpin the ability to integrate information across related experiences in recently acquired memories. Concurrently, EBNs bolster the dynamics of a gradually emerging, long-lasting trace, which proves more resistant to updates. Therefore, our research sheds light on the relationship between the processes governing a memory’s long-term persistence and evolution, and the dynamic nature of the neuronal ensembles in which it is encoded. Moreover, it unveils a potential mechanism underlying the plasticity of days-old memories, grounded in the interplay between developmentally-defined memory networks and the dynamic reorganization of memory ensembles over time.

---

**Fig. S1.**
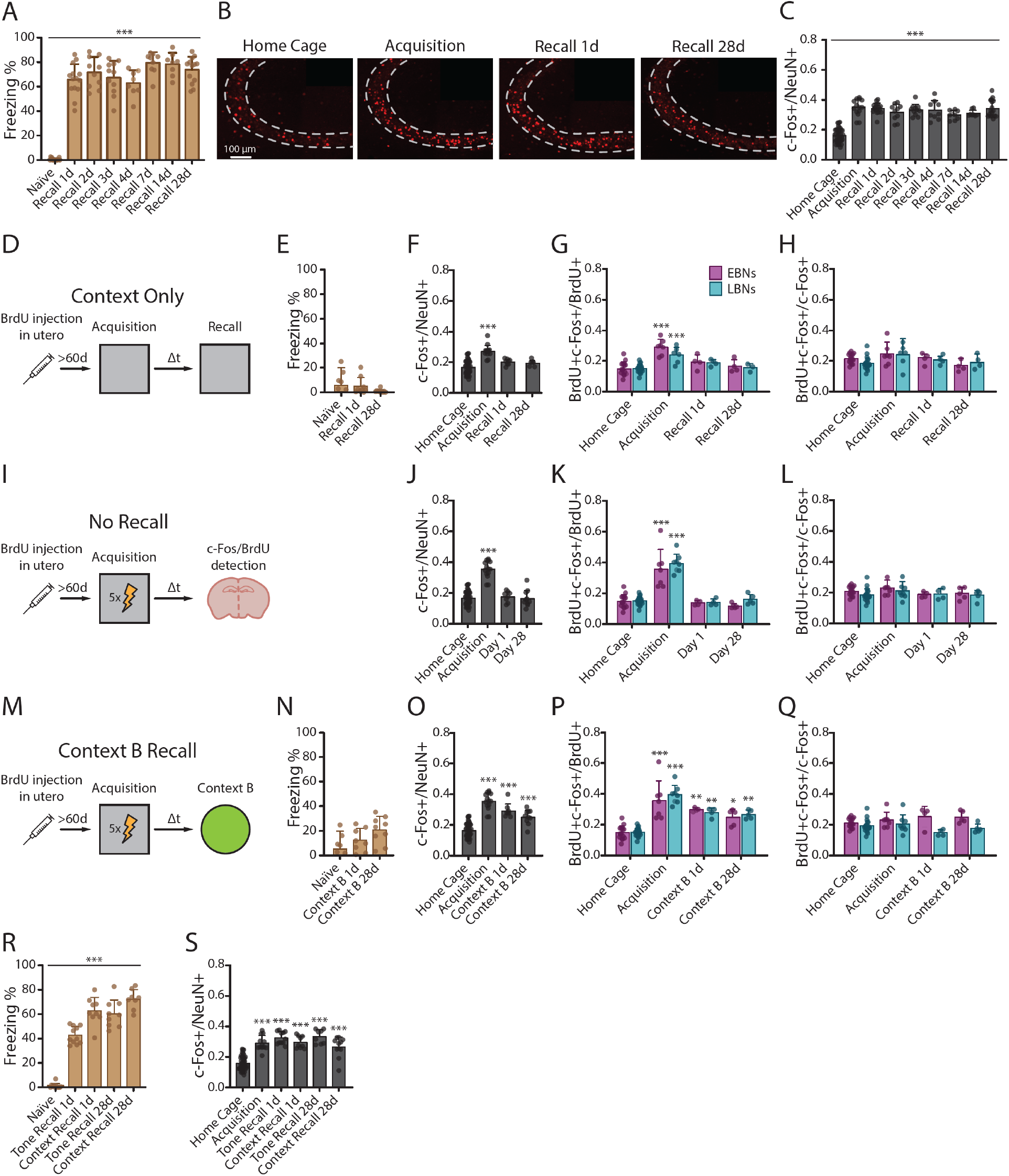
Distinct recruitment dynamics for EBNs and LBNs are restricted to the processing of specific fear-association memories. **(A)** Percentage of time spent freezing (y-axis) upon the exploration of the Training Context before in naïve animals, or upon cFC Recall. Each dot is a different mouse. Bars represent the mean, error bars the SD. cFC Acquisition led to the encoding of a long-lasting fear memory that mice could express at Recall up to 28 days after Acquisition (one-way ANOVA, F(7, 82) = 93.92, p<0.0001. Dunnett’s multiple comparisons test to Naïve: p<0.0001 for all groups). At least 8 animals per group, no repeated measures. **(B)** c-Fos (red) immunolabelling of neurons in the pyramidal layer (dashed line) of Hippocampal area CA3. Micrographs depict representative samples from animals processed directly from their Home Cage, and upon the Acquisition and Recall of a cFC memory. The scale bar is 100 µm, images are maximum intensity projections, 30X silicone objective. **(C)** Fraction of CA3 neurons (NeuN+) expressing c-Fos upon cFC Acquisition or Recall (c-Fos+/NeuN+). Each dot is a different mouse. Bars represent the mean, error bars the SD. Almost twice as many CA3 neurons express c-Fos upon the encoding and retrieval of a cFC memory in comparison to Home Cage controls. One-way ANOVA, F(8, 123) = 43.80, p<0.0001. Dunnett’s multiple comparison test to Home Cage: p<0.0001 for each experimental group. At least 8 animals per group, no repeated measures. **(D)** Schematic of the “Context Only” experiment on BrdU pulse-chased animals. **(E)** Percentage of time spent freezing (y axis) upon the exploration of the Training Context before in naïve animals, or upon Context Only Recall. Each dot is a different mouse. Bars represent the mean, error bars the SD. Mice who were trained in the Context Only paradigm did not exhibit freezing at any time during the Recall of the Contextual memory (one-way ANOVA, F(2,21) = 3.301, p>0.05). At least 8 animals per group, no repeated measures. **(F)** Fraction CA3 neurons (NeuN+) expressing c-Fos upon Context Only Acquisition or Recall (c-Fos+/NeuN+). Each dot is a different mouse. Bars represent the mean, error bars the SD. Exposure to a novel context induced an increase in the expression of c-Fos among CA3 neurons, but no significant increase upon Recall 1 or 28 days after. One-way ANOVA, F(3, 60) = 20.83, p<0.0001. Dunnett’s multiple comparisons test to Home Cage: p<0.0001 for Acquisition, p = 0.098 for Recall 1d, p>0.05 for Recall 28d. At least 8 animals per group, no repeated measures. **(G)** Fraction of CA3 EBNs (magenta) or LBNs (cyan) expressing c-Fos (BrdU+ c-Fos+/BrdU+) upon the encoding and retrieval of a Context-Only memory. Each dot is a different mouse. Bars represent the mean, error bars the SD. Both EBNs and LBNs exhibited a significant increase in c-Fos expression upon the first exposure to the neutral context, but no significant increase upon Recall 1 or 28 days after (for EBNs: one-way ANOVA, F(3,27) = 16.81, p<0.0001. Dunnett’s multiple comparisons test to Home Cage c-Fos expression in EBNs: p<0.0001 for Acquisition, p>0.05 for Recall 1d and 28d. For LBNs: one-way ANOVA, F(3,30) = 12.30, p<0.0001. Dunnett’s multiple comparisons test to Home Cage c-Fos expression in LBNs: p<0.0001 for Acquisition, p>0.05 for Recall 1d and 28d). At least 3 animals per group, no repeated measures. **(H)** Prevalence of EBNs or LBNs in CA3 c-Fos+ ensembles (BrdU+ c-Fos+/c-Fos+) supporting the encoding and retrieval of a Context Only memory. Each dot is a different mouse. Bars represent the mean, error bars the SD. EBNs and LBNs abundance in c-Fos ensembles were never significantly different from baseline levels in Home Cage (for EBNs: one-way ANOVA, F(3, 27) = 1.903, p>0.05. For LBNs: one-way ANOVA, F(3, 30) = 2.775, p>0.05). **(I)** Schematic of the “No Recall” experiment on BrdU pulse-chased animals. **(J)** Fraction of CA3 neurons (NeuN+) expressing c-Fos upon the “No Recall” paradigm (c-Fos+/NeuN+). Each dot is a different mouse. Bars represent the mean, error bars the SD. The Acquisition of a cFC memory induced an increase in the expression of c-Fos among CA3 neurons, but no significant increase if animals were tested 1 or 28 days after (one-way ANOVA F(3, 65) = 59.28, p<0.0001. Dunnett’s multiple comparisons test to Home Cage: p<0.0001 for Acquisition, p>0.05 for Recall 1d and 28d). At least 8 animals per group, no repeated measures. **(K)** Fraction of CA3 EBNs (magenta) or LBNs (cyan) expressing c-Fos (BrdU+ c-Fos+/BrdU+) upon the “No Recall” paradigm. Each dot is a different mouse. Bars represent the mean, error bars the SD. When the animals were not subjected to a recall session after having encoded the cFC memory during Acquisition, we could observe no difference in c-Fos expression in isochronic populations compared to controls neither on Day 1 nor Day 28 after memory encoding. For EBNs: one-way ANOVA, F(3, 28) = 18.97, p<0.0001. Dunnett’s multiple comparisons test to Home Cage: p<0.0001 for Acquisition, p>0.05 for Day 1 and Day 28 For LBNs: one-way ANOVA, F(3, 34) = 93.69, p<0.0001. Dunnett’s multiple comparisons test to home cage: p<0.0001 for Acquisition, p>0.05 for Day 1 and Day 28 groups. At least 4 animals per group, no repeated measures. **(L)** Prevalence of EBNs or LBNs in CA3 c-Fos+ ensembles (BrdU+ c-Fos+/c-Fos+) upon the “No Recall“ paradigm. Each dot is a different mouse. Bars represent the mean, error bars the SD. EBNs and LBNs abundance in c-Fos ensembles were never significantly different than baseline levels in Home Cage (for EBNs: one-way ANOVA, F(3, 28) = 1.170, p>0.05. For LBNs: one-way ANOVA, F(3, 33) = 0.5747, p>0.05). **(M)** Schematic of the “Context B Recall” experiment on BrdU pulse-chased animals. **(N)** Mice who were trained in the Context B Recall paradigm did not exhibit significant levels of freezing at any time during Context B exposure on Day 1 or 28 (one-way ANOVA, F(2,19) = 2.511, p>0.05). At least 8 animals per group, no repeated measures. **(O)** Fraction CA3 neurons (NeuN+) expressing c-Fos upon Context B Recall paradigm (c-Fos+/NeuN+). Each dot is a different mouse. Bars represent the mean, error bars the SD. Exposure to the novel Context B induced increased expression of c-Fos in CA3 neurons at any time after cFC Acquisition, consistent with the fact that on each Context B exposure day, animals were naïve to the exploration of this context (see Fig. S1F). One-way ANOVA F(3, 65) = 59.51, p<0.0001. Dunnett’s multiple comparisons test to Home Cage: p<0.0001 for Acquisition, Context 1d and Context B 28d. At least 8 animals per group, no repeated measures. **(P)** Fraction of CA3 EBNs (magenta) or LBNs (cyan) expressing c-Fos (BrdU+ c-Fos+/BrdU+) upon the Context B Recall paradigm. Each dot is a different mouse. Bars represent the mean, error bars the SD. Exposure to the novel Context B induced increased expression of c-Fos in both EBNs and LBNs at any time after cFC Acquisition. For EBNs: one-way ANOVA, F(3, 28) = 16.09, p<0.0001. Dunnett’s multiple comparisons test to Home Cage: p<0.0001 for Acquisition, p=0.0022 for Context B 1d, and p=0.0299 for Context B 28d. For LBNs: one-way ANOVA, F (3, 34) = 90.05, p<0.0001. Dunnett’s multiple comparisons test to Home Cage: p = 0.0039 for Context B 1d and p = 0.0087 for the Context B 28d groups. At least 4 animals per group, no repeated measures. **(Q)** Prevalence of EBNs or LBNs in CA3 c-Fos+ ensembles (BrdU+ c-Fos+/c-Fos+) upon the Context B Recall paradigm. Each dot is a different mouse. Bars represent the mean, error bars the SD. EBNs and LBNs abundance in c-Fos ensembles were never significantly different than baseline levels in Home Cage (for EBNs: one-way ANOVA, F(3, 28) = 1.595, p>0.05. For LBNs: one-way ANOVA, F(3, 33) = 2.010, p>0.05). **(R)** Percentage of time spent freezing (y-axis) in naïve animals, or upon tFC Acquisition, Tone, and Context Recall. Each dot is a different mouse. Bars represent the mean, error bars the SD. tFC Acquisition leads to the encoding of a long-lasting fear memory that could be expressed by either the presentation of the Context-CS (Context Recall) or the Tone-CS (Tone Recall) up to 28 days after encoding (one-way ANOVA, F(4, 43) = 113.8, p<0.0001. Dunnett’s multiple comparisons test to Naïve animals: p<0.0001 for all groups) At least 8 animals per group, no repeated measures. **(S)** Fraction of CA3 neurons (NeuN+) expressing c-Fos upon the tFC paradigm (c-Fos+/NeuN+). Each dot is a different mouse. Bars represent the mean, error bars the SD. Almost twice as many CA3 neurons expressed c-Fos upon Acquisition and Recall of a tFC memory, independent from the sensory modality used to retrieve the memory at Recall, than the Home Cage controls. One-way ANOVA, F(5, 86) = 41.61, p<0.0001. Dunnett’s multiple comparisons test to Home Cage: p<0.0001 for each experimental group. At least 8 animals per group, no repeated measures.

**Fig. S2.**
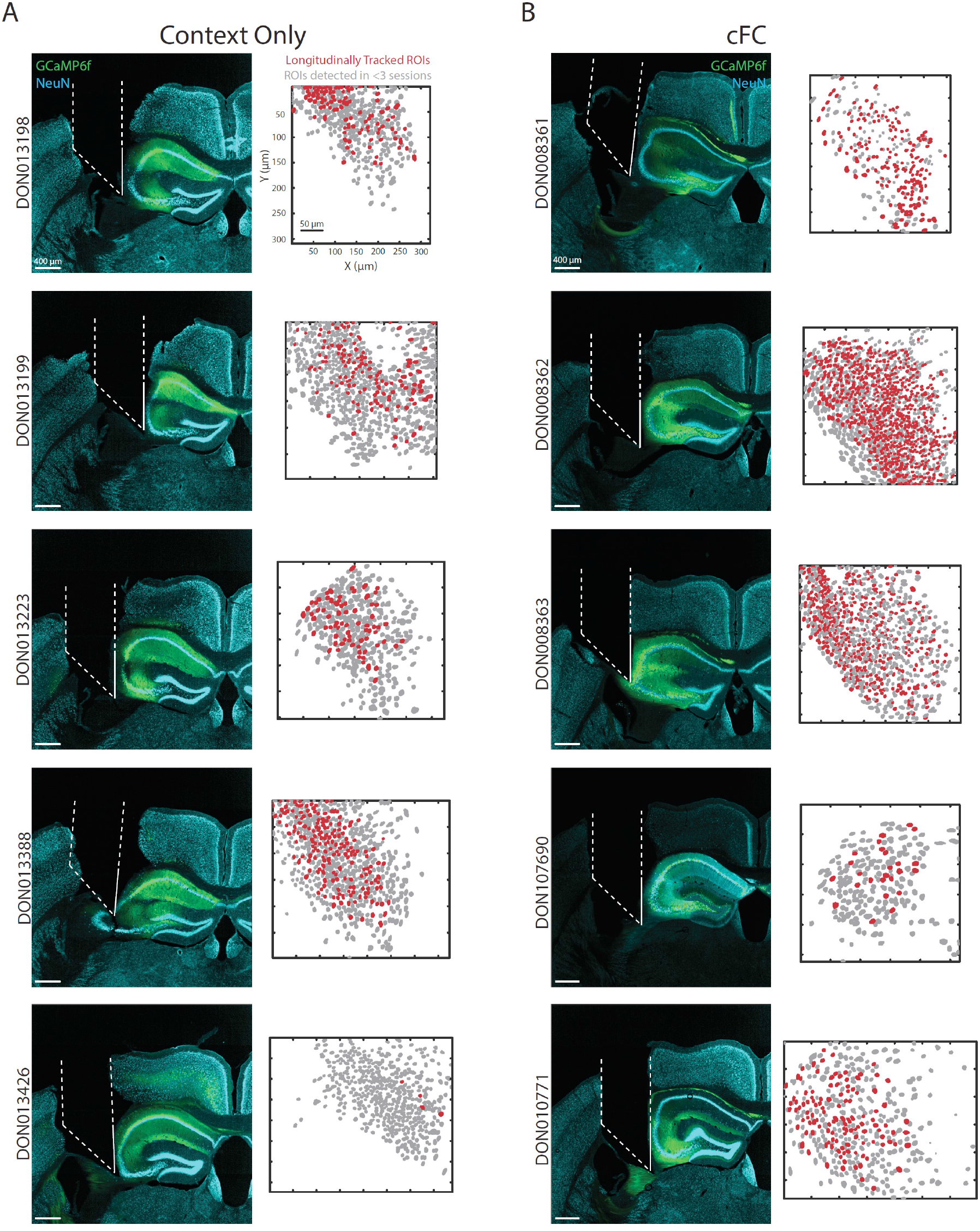
Histological analysis of prism positioning in animals implanted with an endoscope for calcium imaging. (**A**, **B**) Left: representative histology images revealing the position of the endomicroscope (dashed white outline) and the prism imaging face (solid white outline) in relation to the hippocampal formation for each animal of the Context Only (A) or cFC (B) cohorts. GCaMP6f expressing neurons in green, NeuN expressing neurons in light blue. Images are single focal planes acquired with an Olympus Spinning Disk microscope (4X objective). The scale bar is 400 µm. Right: distribution of the recorded Regions of Interest (ROIs) computed to identify neurons that could be reliably tracked across all sessions (red) and neurons that could only be detected in 1 or 2 sessions (dark grey). The scale bar is 50 µm.

**Fig. S3.**
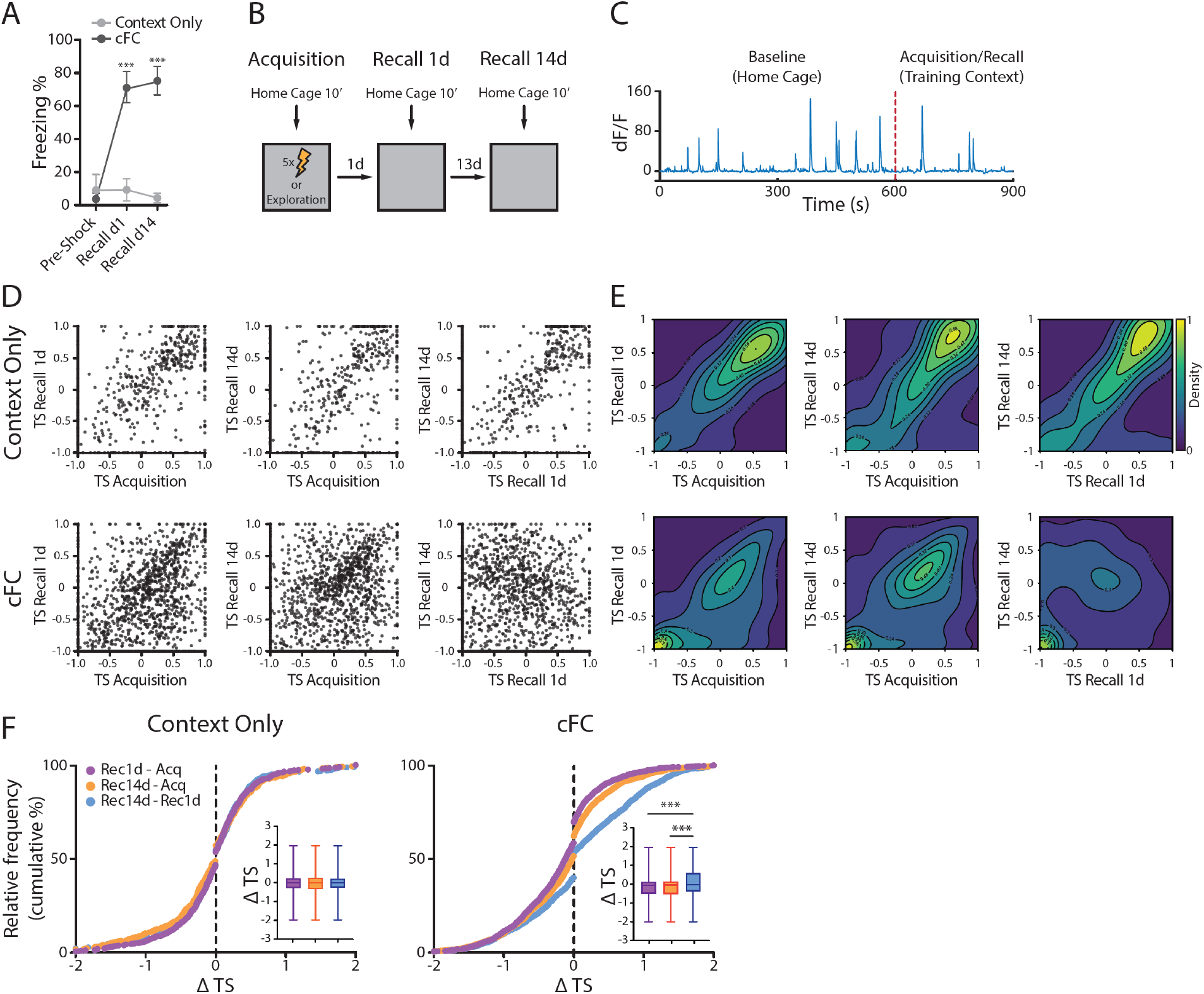
Representation of Training Context orthogonalizes on distinct neuronal subpopulations upon the 1d and 14d Recall of a contextual Fear Memory. **(A)** Percentage of time spent freezing (y-axis) upon the exploration of the Training Context during the initial portion of the Acquisition session (pre-Shock) and upon Recall. Mice trained in the cFC task (N = 5, dark grey) undergoing 2 recall sessions at 1- and 14-days delay from Acquisition, exhibit freezing responses at Recall that are significantly different than mice who never experience a US in the training Context (“Context Only” controls, N = 5 mice, light grey), and significantly higher than their freezing pre-shock during the Acquisition session. Each dot is the average ± SD across mice of a specific cohort on a specific experimental day. Repeated Measures two-way ANOVA performed to analyse the effect of Training Protocol and Experimental Session on the freezing of animals from the cFC cohort revealed that there was a statistically significant interaction between the two variables (F(2,16) = 76.83, p <0.0001). Simple main effects analysis showed that the Training Protocol and Experimental Session both had a statistically significant effect on freezing (F(1,8) = 248, p<0.0001; and (F(1,8) = 66.86, p<0.0001, respectively). A repeated measure one-way ANOVA with Geisser-Greenhouse correction on the cFC cohort revealed a significant difference across experimental sessions (F(1.95,7.80) = 151.1, p<0.0001) but not across individuals (F(4, 8) = 1.536, p>0.05). Dunnett’s multiple comparisons test to pre-shock freezing: p = 0.0003 and p = 0.0001 for Recall 1d and Recall 14d, respectively. A repeated measure one-way ANOVA with Geisser-Greenhouse correction on the Context Only cohort revealed no significant difference across experimental sessions (F(1.156, 4.623) = 0.6469, p>0.05) nor individuals (F(4, 8) = 0.4545, p>0.05). **(B)** Schematic of the imaging experiment. In each experimental session, CA3 activity was first recorded for 10 minutes in the Home Cage to assess the baseline firing rate of recorded neurons. Then the animal was transferred into the Training Context for the cFC/Context Only Acquisition or Recall. Individual animals were imaged longitudinally during Acquisition, Recall 1d, and Recall 14d. **(C)** Example fluorescence trace of one recorded neuron during one experimental session. The Red dashed line demarcates the start of the recording in the Training Context. **(D)** Tuning score for individual neurons across pairs of sessions for the Context Only (top row, N = 503 neurons across 5 animals) and cFC (bottom row, N = 1444 neurons across 5 animals) cohorts. Each scatter plot represents one comparison between two sessions, either between the Acquisition and the Recall 1d or 14d sessions (left and centre panels, respectively), or between the two Recall sessions (right panels). Each dot represents a longitudinally-tracked neuron. Most neurons exhibit preserved tuning between sessions, as indicated by the higher density of data points along the diagonal, for every comparison except the one for cFC Recall 1d Vs Recall 14d. Neurons aligned to the axis represent neurons whose activity is exclusive to one environment, either Home Cage or Training Context. **(E)** 2D kernel density estimation of scatterplots from Fig. S3D. Colours indicate the normalized value of the underlying probability density function estimated from the distribution of data points from Fig. S3D. Contour lines connect areas with similar probability density, indicated adjacently to the contour. **(F)** Cumulative distribution across the population for individual neuron’s differences in tuning scores between pairs of sessions (1: Acquisition vs. Recall 1d, in purple; 2: Acquisition vs. Recall 14d, in orange; 3: Recall 1d vs. Recall 14d, in blue). While in the Context Only cohort (left), the distribution of ΔTS is similar for every comparison (Kolmogorov-Smirnov test: 2 vs. 1: D = 0.02, p>0.05; 3Vs1: D = 0.017, p>0.05; 3 vs. 2: D = 0.025, p>0.05), in the cFC cohort (right) the distribution of ΔTS across the two Recall sessions was significantly different from the others (Kolmogorov-Smirnov test. 2 vs. 1: D = 0.04, p>0.05; 3 vs. 1: D = 0.134, p<0.0001; 3 vs. 2: D = 0.114, p<0.0001). Insets: box plots indicating the distribution of ΔTS across experimental sessions for animals from the Context Only (left) and cFC (right) cohorts. Individual animals (5 per cohort) are pooled. Repeated measures one-way ANOVA revealed no significant difference in the distribution of ΔTS for each comparison in the Context Only cohort (F(1.262, 633.5) = 1.064, p>0.05), and a significant difference across comparisons in the cFC cohort (F(1.173, 1693) = 81.93, p<0.0001. Tukey’s multiple comparisons test: p<0.0001 for each pair of comparisons).

**Figure S4.**
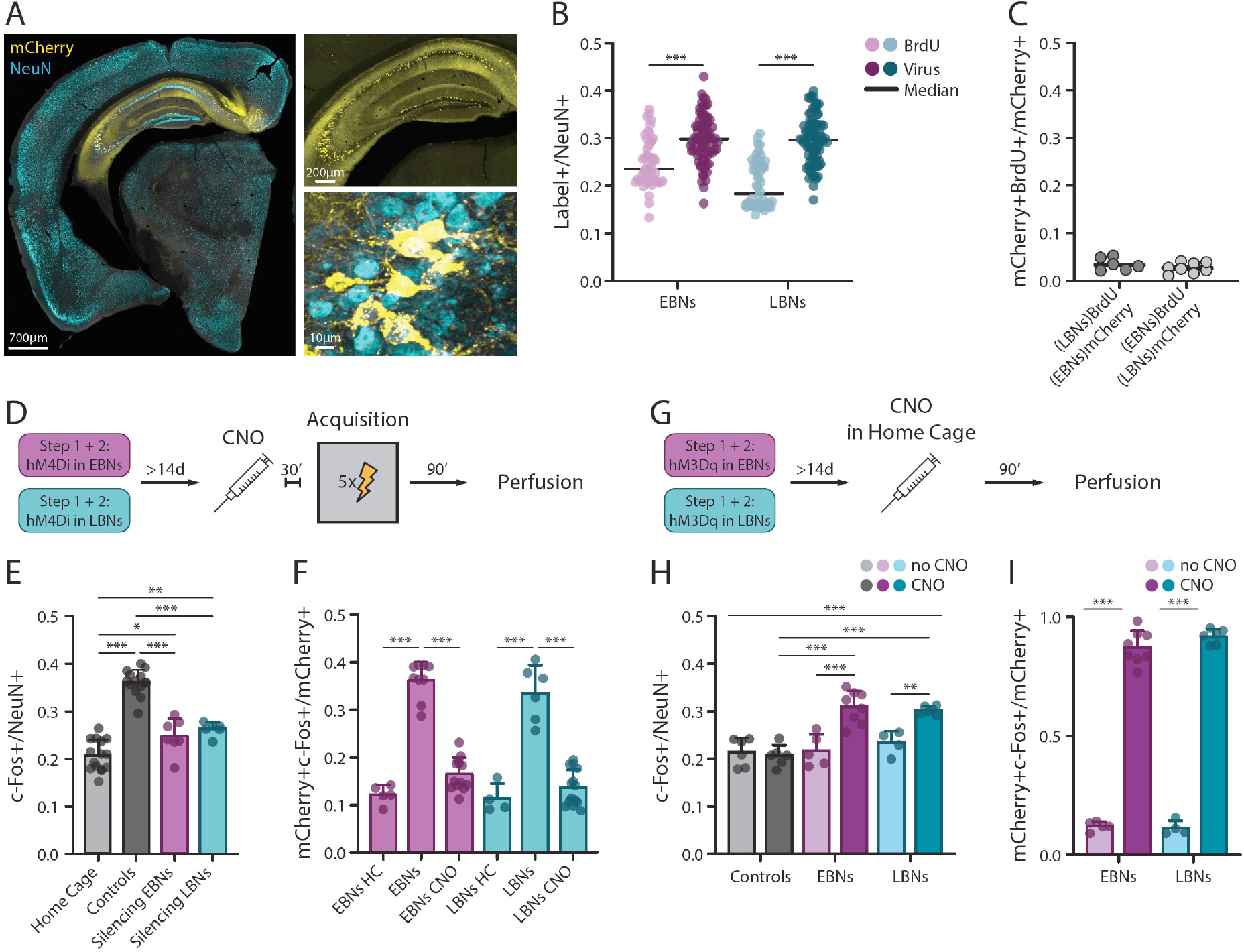
Chemogenetic manipulation of isochronic neurons’ recruitment in c-Fos+ ensembles. **(A)** mCherry (yellow) and NeuN (cyan) immunolabelling in the brain of LBNs-targeted, DREADD-expressing mice. Left: maximum intensity projection image of a whole hemisphere acquired with an Olympus Spinning Disk, 10X air objective. Right: maximum intensity projection image of the whole hippocampus (upper panel), and zoom in towards CA3 neurons (lower panel), acquired with an Olympus Spinning Disk, 30X silicone objective. **(B)** Fraction of the CA3 network labelled by the pulse-chase (BrdU+) or all-viral (mCherry+) method to target isochronic subpopulations during development. EBNs (magenta) were targeted by injection of BrdU (light shade) or an AAV virus (dark shade) at E12, while LBNs (cyan) were targeted by injection of BrdU (light shade) or an AAV virus (dark shade) at E15. While roughly the same proportion of EBNs and LBNs CA3 neurons are labelled within the same method, we can observe a consistently larger fraction of the network being labelled via the all-viral approach (for EBNs targeting: two-tailed unpaired t-test, t(120) = 5.411, p<0.0001. For LBNs targeting: two-tailed unpaired t-test, t(120) = 10.33, p<0.0001). This could be a consequence of a longer temporal window for isochronic neuron labelling characterizing the all-viral method in comparison to the BrdU pulse-chase method (*67*). At least 50 animals per group. **(C)** Fraction of double-positive BrdU+/mCherry+ neurons when both subpopulations are targeted in parallel in the same animal via the pulse-chase and the all-viral method (as in Fig. 3A). The low fraction of double-labelled neurons reveals that the all-viral method specifically labels the isochronic population it is targeted towards, with minimal contamination by the non-targeted subpopulation. **(D)** Schematic of the experiment for the silencing of EBNs and LBNs upon cFC Acquisition. **(E)** Fraction of CA3 neurons (NeuN+) expressing c-Fos in Home Cage animals (N = 9), upon cFC Acquisition in Controls (*i.e.* animals where ENBs or LBNs expressed DREADD, but no ligand CNO was injected. N = 15 mice), and upon DREADD-mediated silencing of EBNs (N = 13) or LBNs (N = 15). Each dot is a different mouse. Bars represent the mean, error bars the SD. Silencing EBNs or LBNs upon Acquisition, a time when both isochronic populations exhibit an increased expression of c-Fos, significantly reduced the size of the c-Fos+ ensemble supporting memory encoding (one-way ANOVA, F(3, 38) = 69.04, p<0.0001. Tukey’s multiple comparisons test: p<0.0001 for Home Cage vs. Controls; p = 0.0238 for Home Cage vs. EBNs Silencing; p = 0.0028 for Home Cage vs. Silencing LBNs; p<0.0001 for Controls vs. Silencing EBNs; p<0.0001 for Controls vs. Silencing LBNs; p>0.05 for Silencing EBNs vs. Silencing LBNs). **(F)** c-Fos expression in isochronic-targeted, DREADD (hM4Di)-expressing CA3 neurons in Home Cage animals (EBN-targeted: N = 5; LBNs-targeted: N = 4), upon cFC Acquisition without CNO Injection (EBNs-targeted: N = 9; LBNs-targeted: N = 6), and upon cFC Acquisition with CNO injection (EBN-targeted, CNO injected: N = 13; LBN-targeted, CNO injected: N = 14). Each dot is a different mouse. Bars represent the mean, error bars the SD. Silencing isochronic neurons at Acquisition prevented the encoding-induced increase in c-Fos expression within the targeted subpopulation, thereby validating the effectiveness of the all-viral isochronic silencing approach. For EBN targeting: one-way ANOVA, F(2,24) = 115, p<0.0001. Tukey’s multiple comparisons test: EBNs Home Cage vs. EBNs, p<0.0001; EBNs Home Cage vs. EBNs CNO, p>0.05; EBNs vs. EBNs CNO, p<0.0001. For LBN targeting: one-way ANOVA, F(2,21) = 51.45, p<0.0001. Tukey’s multiple comparisons test: LBNs Home Cage vs. LBNs, p<0.0001; LBNs Home Cage vs. LBNs CNO, p>0.05; LBNs vs. LBNs CNO, p<0.0001. **(G)** Schematic of the experiment for the excitation of EBNs and LBNs upon baseline conditions (Home Cage). Adult (>P60) mice where EBNs (magenta) or LBNs (cyan) had been targeted with the all-viral method described in Fig. 2A for the expression of the excitatory version of the DREADD receptor, were injected with CNO to induce the activation of hM3Dq and consequent activation of isochronic neurons, and housed in their Home Cage until their brain was processed for further analysis. **(H)** Fraction of CA3 neurons (NeuN+) expressing c-Fos in Home Cage animals without CNO injection (N = 6, grey, light shade) or with CNO injection (N = 6, grey, dark shade); EBN-targeted without CNO (N = 5, magenta, light shade) or with CNO (N = 8, magenta, dark shade); and LBN-targeted without CNO (N = 4, cyan, light shade) or with CNO (N = 6, cyan, dark shade). Each dot is a different mouse. Bars represent the mean, error bars the SD. The activation of EBNs or LBNs in the Home Cage significantly increased c-Fos expression in the CA3 network (one-way ANOVA, F(5, 29) = 18.40, p<0.0001. Tukey’s multiple comparisons test indicated significant differences for Controls no CNO vs. EBNs CNO (p<0.0001), Controls CNO vs. EBNs CNO (p<0.0001), Controls no CNO vs. LBNs CNO (p<0.0001), Controls CNO vs. LBNs CNO (p<0.0001), EBNs no CNO vs. EBNs CNO (p<0.0001), LBNs no CNO vs. LBNs CNO (p = 0.0079), EBNs no CNO vs. LBN CNO (p = 0.0003), LBNs no CNO vs. EBNs CNO (p = 0.0015). p>0.05 for the other comparisons. **(I)** c-Fos expression in EBNs of EBN-targeted animals without CNO injection (N = 5) or with CNO injection (N = 8) in their Home Cage; and LBNs of LBN-targeted animals without CNO injection (N = 4) or with CNO injection (N = 6) in their Home Cage. Each dot is a different animal. Bars represent the mean, error bars the SD. The activation of isochronic neurons induced increased expression of c-Fos in the targeted subpopulation, thereby validating the effectiveness of the all-viral isochronic excitation approach. For EBN targeting: unpaired two-tailed t-test, t(11) = 22.96, p<0.0001. For LBN targeting: unpaired two-tailed t-test, t(8) = 40.58, p<0.0001.

**Figure S5.**
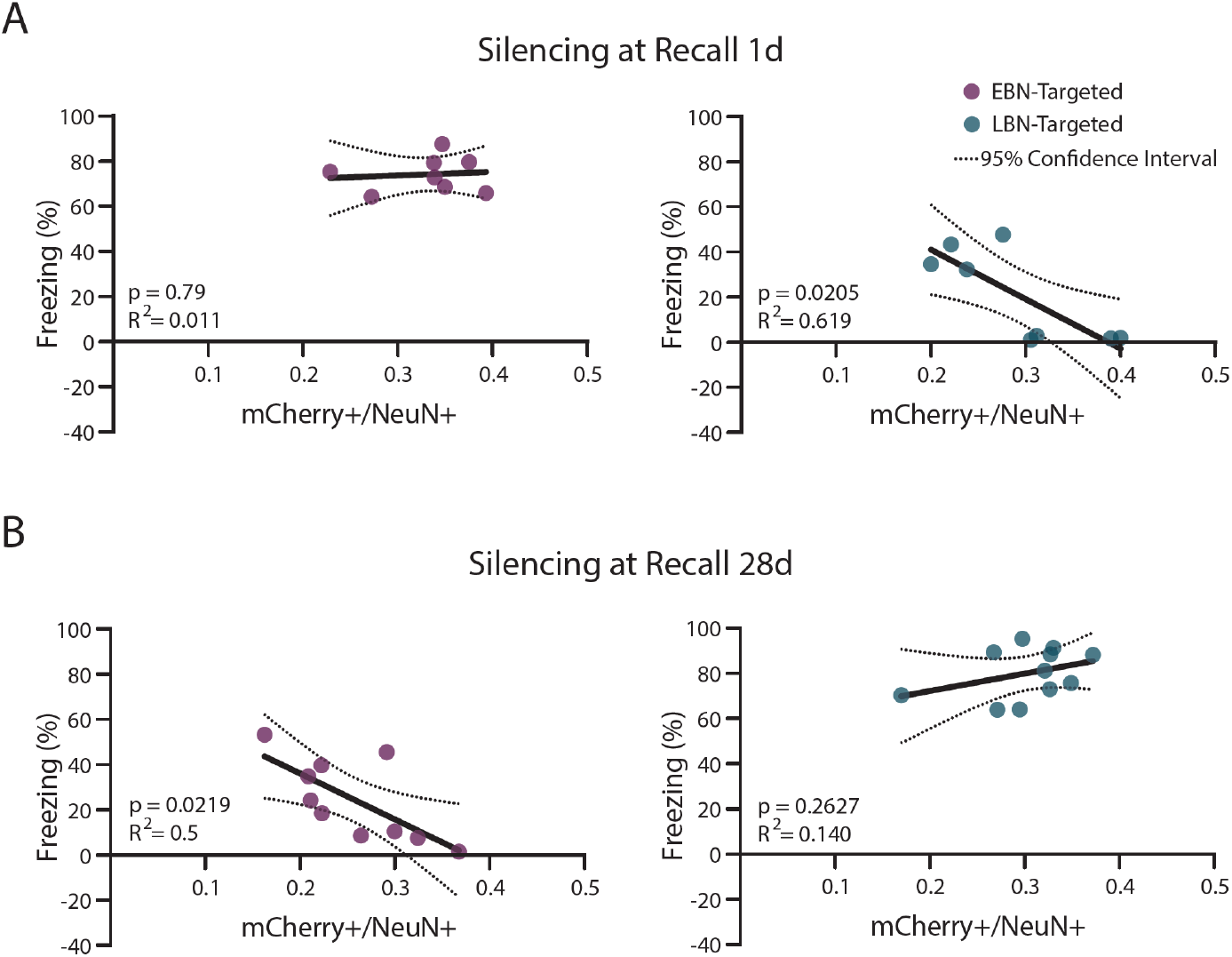
Memory retrieval impairment correlates with subpopulation-specific isochronic silencing at Recall. (**A**, **B**) A simple linear regression was calculated to predict freezing at Recall based on the fraction of the CA3 network that expressed the DREADD receptor hM4Di when this was targeted to EBNs or LBNs. A significant regression was found for (A, right panel) LBN-targeted animals when silencing was performed upon Recall 1d (F(1,6) = 9.759, p = 0.0205), with an R^2^ of 0.619, and for (B, left panel) EBN-targeted animals when silencing was performed upon Recall 28d (F(1,8) = 8.049, p = 0.0219), with an R^2^ of 0.50. No significant regression was found for (A, left panel) EBN-targeted animals when silencing was performed upon Recall 1d (F(1,6) = 0.072, p>0.05) with an R^2^ of 0.011, and for (B, right panel) LBN-targeted animals when silencing was performed upon Recall 28d (F(1,9) = 1.428, p>0.05), with an R^2^ of 0.140. Sample size as in Fig. 2D and 2E. The dotted lines represent 95% confidence interval.

**Figure S6.**
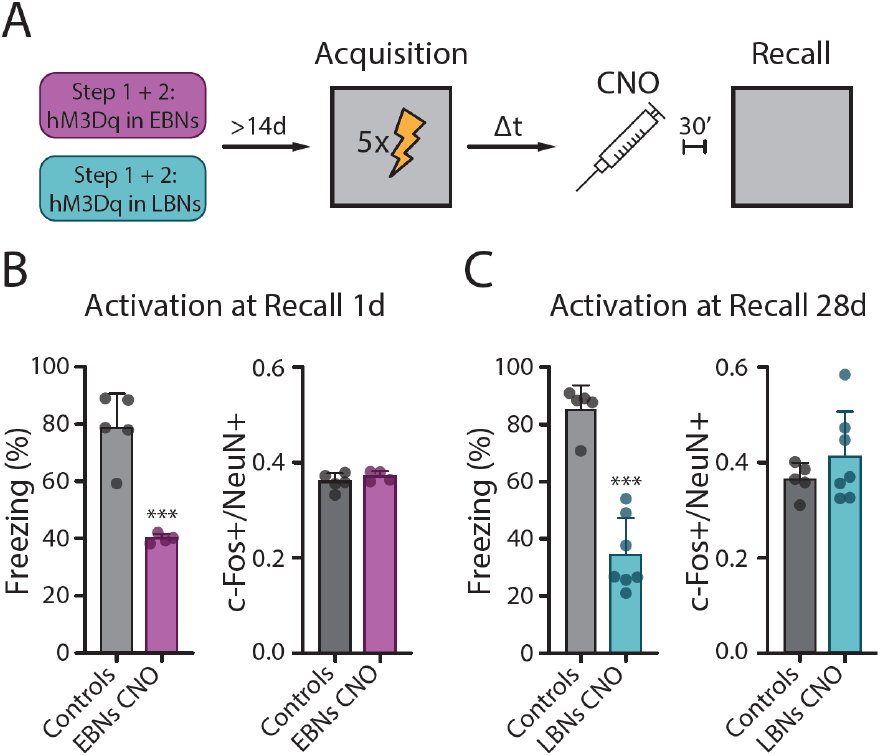
Anachronistic activation of isochronic subpopulations impairs cFC memory retrieval. **(A)** Schematic of the experiment for the activation of EBNs and LBNs at cFC Recall. As in Fig. 2C, with the difference that the activator version of the DREADD receptor hM3Gq was targeted to EBNs and LBNs during Step 2 of the all-viral approach. **(B)** Left: freezing during the exploration of the Training Context at Recall 1d in Control animals (N = 5) and upon DREADD-mediated activation of EBNs (N = 4). Each dot is a different mouse. Bars represent the mean, error bars the SD. Memory retrieval was significantly impaired upon the activation of EBNs at Recall 1d (two-tailed student’s t-test, t(7) = 6.291, p = 0.0004). Right: c-Fos expression in the CA3 network upon the activation of EBNS at Recall 1d. Y-axis: fraction of c-Fos+, NeuN+ neurons normalized by the total number of neurons in the CA3 network (NeuN+). Each dot is a different mouse. Bars represent the mean, error bars the SD. The size of the c-Fos+ ensemble recruited upon memory retrieval did not change upon activation of EBNs (two-tailed student’s t-test, t(7) = 1.122, p>0.05). **(C)** Left: freezing during the exploration of the Training Context at Recall 28d in Control animals (N = 5) and upon DREADD-mediated Activation of LBNs (N = 7). Each dot is a different mouse. Bars represent the mean, error bars the SD. Memory retrieval was significantly impaired upon the activation of LBNs at Recall 28d (Two-tailed student’s t-test, t(10) = 7.766, p<0.0001). Right: c-Fos expression in the CA3 network upon the activation of LBNs at Recall 28d. y axis: fraction of c-Fos+, NeuN+ neurons normalized by the total number of neurons in the CA3 network (NeuN+). Each dot is a different mouse. Bars represent the mean, error bars the SD. The size of the c-Fos+ ensemble recruited upon memory retrieval did not change upon activation of LBNs (two-tailed student’s t-test, t(10) = 1.051, p>0.05).

**Figure S7.**
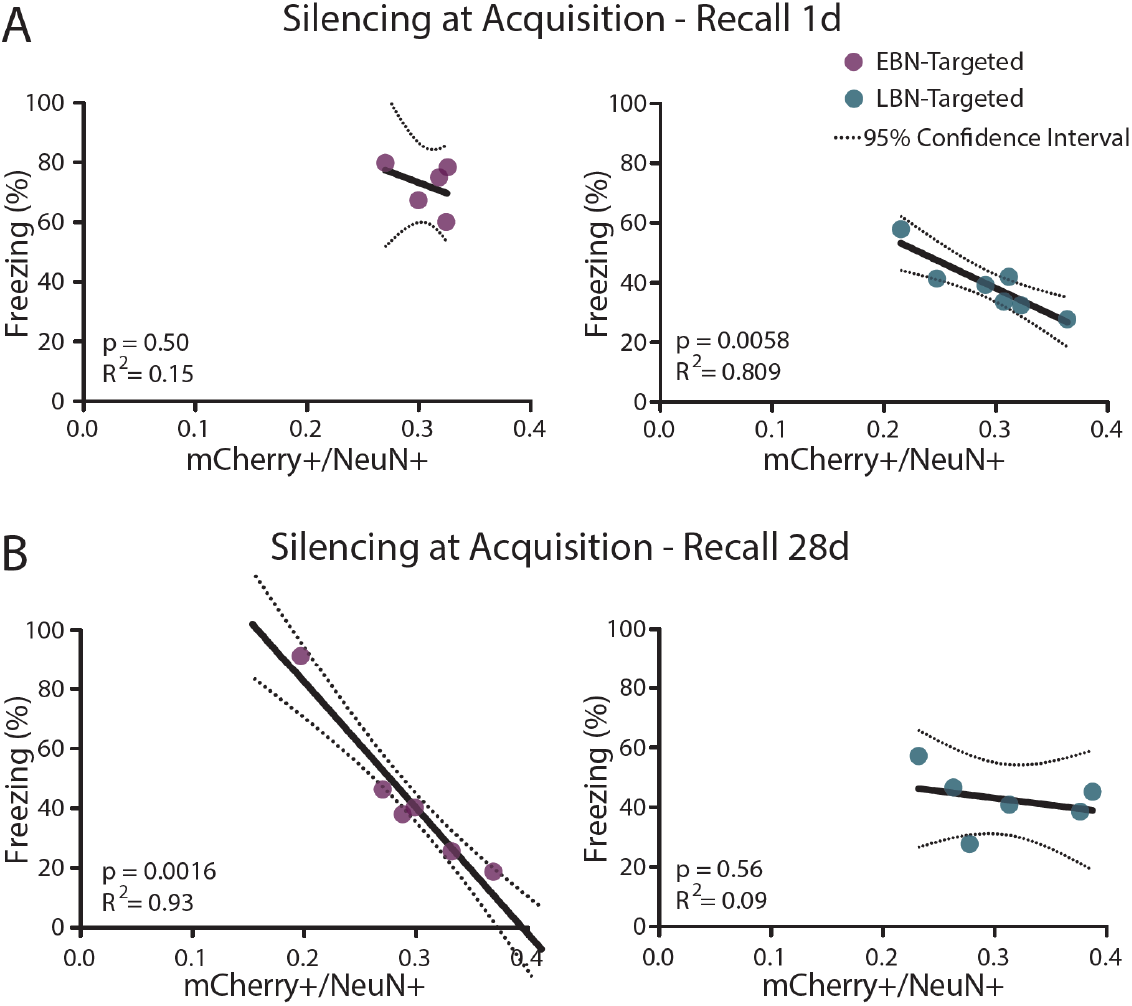
Memory retrieval impairment correlates with subpopulation-specific isochronic silencing at Acquisition. (**A, B**) A simple linear regression was calculated to predict freezing at Recall based on the fraction of the CA3 network that expressed the DREADD receptor hM4Di when this was targeted to EBNs or LBNs, and Silencing was performed at Acquisition. A significant regression was found for (A, right panel) LBN-targeted animals upon Recall 1d (F(1,5) = 21.17, p = 0.0058), with an R^2^ of 0.809, and for (B, left panel) EBN-targeted, DREADD-expressing animals upon Recall 28d (F(1,4) = 57.15, p = 0.0016), with an R^2^ of 0.93. No significant regression was found for (A, left panel) EBN-targeted animals upon Recall 1d (F(1,3) = 0.56, p>0.05), with an R^2^ of 0.15, and for (B, right panel) LBN-targeted animals upon Recall 28d (F(1,4) = 0.39, p>0.05), with an R^2^ of 0.09. Samples size as in Fig. 3, G and H. The dotted lines represent 95% confidence interval.

**Fig. S8.**
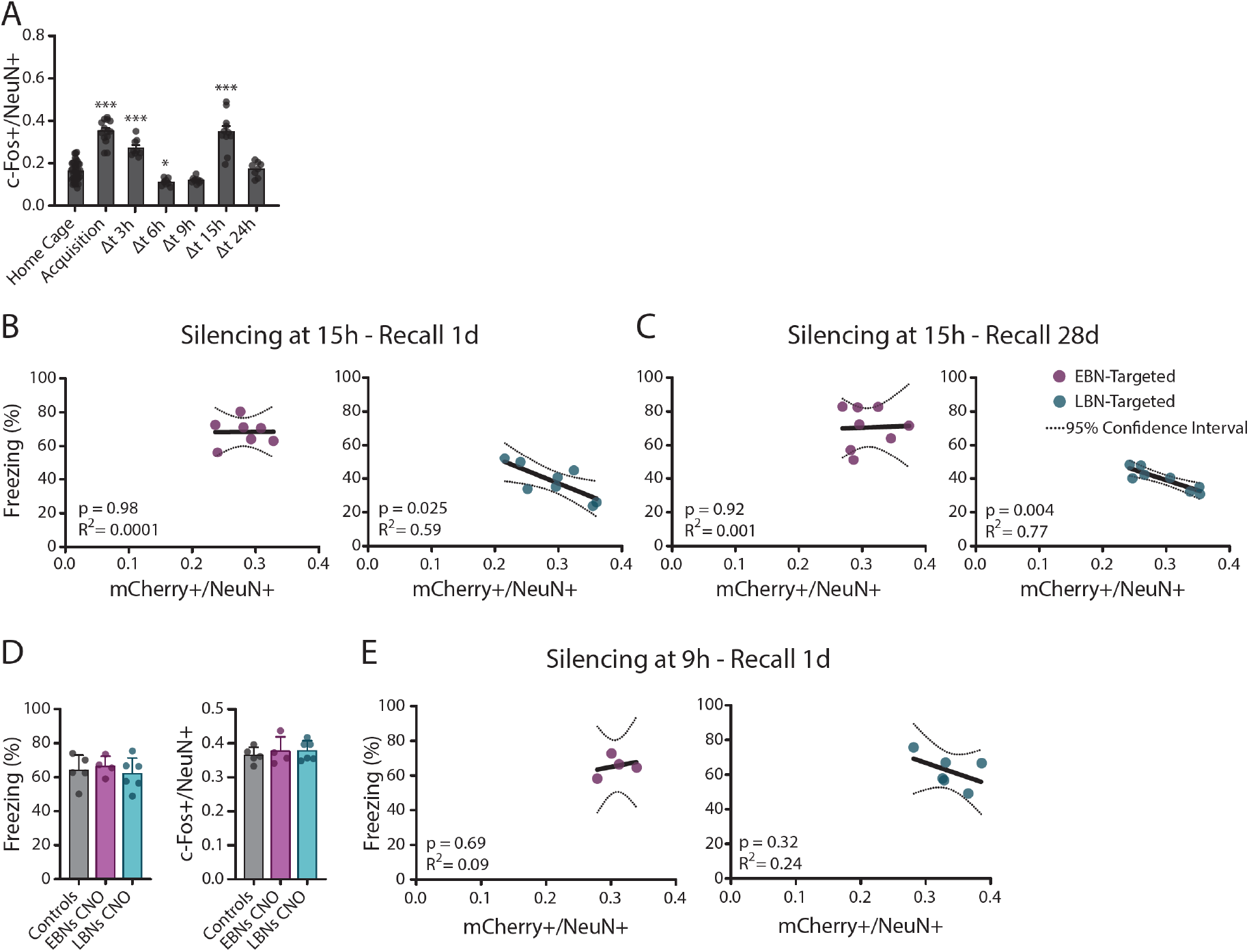
Memory retrieval impairment correlates with LBN silencing during a specific window of consolidation. (**A**) Fraction CA3 neurons (NeuN+) expressing c-Fos in the hours following the encoding of a cFC memory. Each dot is an animal. Bars represent the mean, error bars the SD. c-Fos expression in CA3 neurons exhibited a peak that started at Acquisition and remained elevated 3 hours after, but was back to baseline by 6h. A second peak was centred around 15h after memory encoding (one-way ANOVA, F(6, 89) = 52.99, p<0.0001. Dunnett’s multiple comparisons test to Home Cage LBNs: p<0.0001 for Acquisition, Δt 3h, and Δt 15h; p = 0.49 for Δt 6h; p>0.05 for Δt = 9h, and 24h). At least 8 animals per group, no repeated measures. (**B**, **C**) A simple linear regression was calculated to predict freezing at Recall based on the fraction of the CA3 network that expressed the DREADD receptor hM4Di when this was targeted to EBNs or LBNs, and silencing was performed during the second wave of c-Fos expression in Home Cage (*i.e.* 15 hours post-Acquisition, CNO injected at 13.5 hours). A significant regression was found for (B, right panel) LBN-targeted animals upon Recall 1d (F(1,6) = 8,757, p = 0.025), with an R^2^ of 0.59, and for (C, right panel) LBN-targeted animals upon Recall 28d (F(1,6) = 20.56, p = 0.004), with an R^2^ of 0.77. No significant regression was found for (B, left panel) EBN-targeted animals upon Recall 1d (F(1,5) = 0.0006, p>0.05), with an R^2^ of 0.0001, and for (C, left panel) EBN-targeted animals upon Recall 28d (F(1,6) = 0.01, p>0.05), with an R^2^ of 0.001. Samples size as in Fig. 3, L and M. The dotted lines represent 95% confidence interval. **(D)** Left: freezing during the exploration of the Training Context at Recall 1d in Control animals (N = 5) and upon DREADD-mediated silencing of EBNs (N = 7) or LBNs (N = 8) at a time during consolidation when neither EBNs nor LBNs express high levels of c-Fos (*i.e.* 9 hours post-Acquisition, CNO injection at 7.5h). Each dot is a different mouse. Bars represent the mean, error bars the SD. Memory retrieval was not significantly altered by isochronic silencing (one-way ANOVA, F(2, 12) = 0.3221, p>0.05). Right: c-Fos expression in the CA3 network at Recall 1d upon the activation of EBNs and LBNS around 9 hours after Acquisition. Y-axis: fraction of c-Fos+, NeuN+ neurons normalized by the total number of neurons in the CA3 network (NeuN+). Each dot is a different mouse. Bars represent the mean, error bars the SD. The size of the c-Fos+ ensemble recruited upon memory retrieval was not significantly different upon isochronic silencing (one-way ANOVA, F(2, 12) = 0.3059, p>0.05). **(E)** A simple linear regression was calculated to predict freezing at Recall based on the fraction of the CA3 network that expressed the DREADD receptor hM4Di when this was targeted to EBNs or LBNs, and silencing was performed between the first and second wave of c-Fos expression in Home Cage (*i.e.* 9 hours post-Acquisition, CNO injection at 7.5h). No significant regression was found for either (E, left panel) EBN-targeted animals upon Recall 1d (F(1,2) = 0.203, p>0.05), with an R^2^ of 0.09, or for (E, right panel) LBN-targeted animals upon Recall 1d (F(1,4) = 1.283, p>0.05), with an R^2^ of 0.24. Sample size as in Fig. S8D. The dotted lines represent 95% confidence interval.

**Fig. S9.**
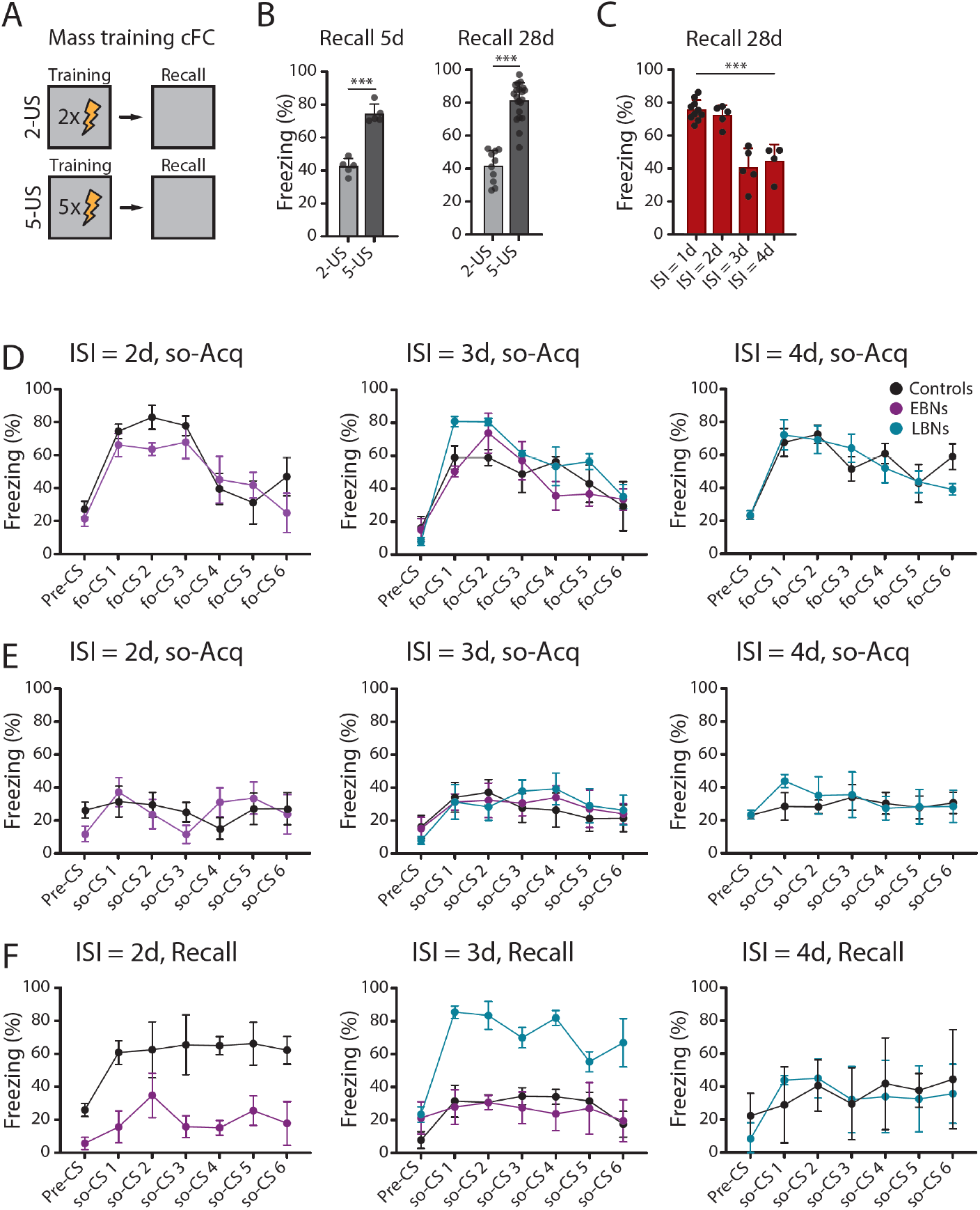
Integration of learning episodes modulated by EBNs and LBNs recruitment in c-Fos+ ensembles. **(A)** Schematic of the Mass training cFC task upon the presentation of a train of 2-US (top row) or 5-US (bottom row) during Training. **(B)** Freezing for animals exploring the Training context during Recall on day 5 (left) or 28 (right) after Acquisition of the Mass training cFC task. Each dot is a different mouse. Left: training in the 2-US (N = 5 mice) and 5-US (N = 5 mice) regimes led to the encoding of a fear memory whose strength scaled with the numbers of shocks experienced during Training (unpaired t-test, t(8) = 9.360, p<0.0001), with a weak-fear memory established upon the presentation of 2 US, and a strong-fear memory established upon the presentation of 5 US. These differences persisted up to 28 days after Acquisition (right. N = 10 for animals trained in the 2-US regime; N = 21 for animals trained in the 5-US regime; unpaired t-test, t(29) = 9.731, p<0.0001). **(C)** Freezing for animals exploring the Training Context during the Recall 28d session of the Spaced cFC task. Each dot is a different animal. Animals who trained in the spaced cFC protocol (N = 5 animals for each group) could combine the two experiences for the production of a strong-fear memory if the Re-Training session was performed with an ISI≤2 days from Training, and this strong-fear memory was maintained for Recall 28d, while animals trained with ISI≥3 days delay maintained a weak-fear memory at Recall 28d (One-way ANOVA, F(3, 15) = 20.58. Tukey’s multiple comparisons test: p = 0.0001 for ISI = 1d vs. ISI = 3d; p = 0.0007 for ISI = 1d vs. ISI = 4d; p = 0.0002 for ISI=2d vs. ISI=3d; p=0.0011 for ISI = 2d vs. ISI = 4d. p>0.05 for all other comparisons). **(D)** Freezing upon the individual presentations of the fo-CS (fo-CS 1 to 6) during so-Acq session upon the activation of isochronic neurons. Samples size as in Fig. 4J-L, mean ± SD. Left: for activation of EBNs at ISI = 2, repeated measures two-way ANOVA with Geisser-Greenhouse correction considering Time (*i.e.* fo-CS presentations from 1 to 6) and Training Protocol (*i.e.* EBN activation vs. Controls) identifies no combined effect of Time x Protocol (F(6, 42) = 1.105, p>0.05) or Protocol (F(1, 7) = 0.9809, p>0.05), but a significant difference when considering Time (F(3.163, 22.14) = 13.26, p<0.0001) individually. Centre: for activation of EBNs or LBNs at ISI = 3, repeated measures two-way ANOVA with Geisser-Greenhouse correction considering Time (*i.e.* fo-CS presentations from 1 to 6) and Training Protocol (*i.e.* EBN or LBN activation vs. Controls) identifies no combined effect of Time x Protocol (F (12, 60) = 1.067, p>0.05), or Protocol (F(2, 10) = 2.608, p>0.05), but a significant difference when considering Time (F(4.193, 41.93) = 16.88, p<0.0001) individually. Right: for activation of LBNs at ISI = 4, repeated measures two-way ANOVA with Geisser-Greenhouse correction considering Time (*i.e.* fo-CS presentations from 1 to 6) and Training Protocol (*i.e.* LBN activation vs. Controls) identifies no combined effect of Time x Protocol (F(6, 42) = 1.843, p>0.05), or Protocol (F(1, 7) = 0.05678, p>0.05), but a significant difference when considering Time (F(3.530, 24.71) = 19.00, p<0.0001) individually. **(E)** Freezing upon the individual presentations of the so-CS (so-CS 1 to 6) during so-Acq session upon the activation of isochronic neurons. Samples size as in Fig. 4J-L, mean ± SD. Left: for activation of EBNs at ISI = 2, repeated measures two-way ANOVA with Geisser-Greenhouse correction considering Time (*i.e.* so-CS presentations from 1 to 6) and Training Protocol (*i.e.* EBN activation vs. Controls) identifies no combined effect of Time x Protocol (F(6, 42) = 0.7414, p>0.05), nor Time (F (2.971, 20.80) = 0.8227, p>0.05) or Protocol (F(1, 7) = 1.677e-006, p>0.05) individually. Centre: for activation of EBNs or LBNs at ISI = 3, repeated measures two-way ANOVA with Geisser-Greenhouse correction considering Time (*i.e.* so-CS presentations from 1 to 6) and Training Protocol (*i.e.* EBN or LBN activation vs. Controls) identifies no significant combined effect of Time x Protocol (F(12, 60) = 0.2741 p>0.05), nor of Time (F(3.853, 38.53) = 2.057, p>0.05) or Protocol (F(2, 10) = 0.3271, p>0.05) individually. Right: for activation of LBNs at ISI = 4, repeated measures two-way ANOVA with Geisser-Greenhouse correction considering Time (*i.e.* so-CS presentations from 1 to 6) and Training Protocol (*i.e.* LBN activation vs. Controls) identifies no combined effect of Time x Protocol (F(6, 42) = 0.3453, p>0.05), nor Time (F(3.358, 23.51) = 0.6367, p>0.05) or Protocol (F(1, 7) = 0.4848, p>0.05) individually. **(F)** Freezing upon the presentation of the so-CS (so-CS 1 to 6) during so-tFC Recall session in animals where isochronic neurons had been activated during the so-Acq session. Samples size as in Fig. 4J-L, mean ± SD. Left: for activation of EBNs at ISI = 2, repeated measures two-way ANOVA with Geisser-Greenhouse correction considering Time (*i.e.* so-CS presentations from 1 to 6) and Training Protocol (*i.e.* EBN activation vs. Controls) identifies no combined effect of Time x Protocol (F(6, 42) = 0.7530, p>0.05), or Time considered individually (F (2.829, 19.81) = 2.637, p>0.05), but a significant difference given by the Protocol (F(1,7) = 16.71, p = 0.0046). Centre: for activation of EBNs or LBNs at ISI = 3, repeated measures two-way ANOVA with Geisser-Greenhouse correction considering Time (*i.e.* so-CS presentations from 1 to 6) and Training Protocol (*i.e.* EBN or LBN activation vs. Controls) identifies a significant combined effect of Time x Protocol (F(12, 60) = 2.212, p = 0.022), and of Time (F(3.755, 37.55) = 7.363, p = 0.0002), and Protocol (F(2, 10) = 20.53, p=0.0003) individually. Dunnett’s multiple comparison test to Controls reveals a statistical difference for LBNs, but not EBNs, Activation. Right: for activation of LBNs at ISI = 4, repeated measures two-way ANOVA with Geisser-Greenhouse correction considering Time (*i.e.* so-CS presentations from 1 to 6) and Training Protocol (*i.e.* LBN activation vs. Controls) identifies no combined effect of Time x Protocol (F(6, 42) = 0.9903, p>0.05), nor Time (F(3.533, 24.73) = 1.375, p>0.05) or Protocol (F(1, 7) = 0.04450, p>0.05) individually.

**Table S1:** Imaging parameters for time-lapse calcium imaging of CA3 neurons. Focus, LED power, Gain, and frame rate for Context Only (N = 5) and cFC (N = 5) mice were set independently for each mouse during the first imaging session and kept constant throughout the experiment.

| Animal ID | Focus<br>(a.u.) | LED Power<br>(a.u.) | Gain<br>(a.u.) | Acquisition rate<br>(Hz) | Group |
| --- | --- | --- | --- | --- | --- |
| **DON013198** | 640 | 0.5 | 3.5 | 25 | Context Only |
| **DON013199** | 440 | 0.5 | 3.6 | 25 | Context Only |
| **DON013223** | 380 | 0.5 | 3.5 | 25 | Context Only |
| **DON013338** | 630 | 0.5 | 3.5 | 25 | Context Only |
| **DON013426** | 380 | 0.4 | 2.0 | 25 | Context Only |
| **DON008361** | 420 | 1.0 | 2.8 | 20 | cFC |
| **DON008362** | 380 | 0.8 | 2.3 | 20 | cFC |
| **DON008363** | 550 | 0.5 | 2.0 | 20 | cFC |
| **DON010769** | 820 | 0.4 | 3.0 | 25 | cFC |
| **DON010771** | 500 | 0.4 | 3.0 | 25 | cFC |

## Materials and Methods

All procedures related to animal housing, surgery, behavioural experiments, and euthanasia were conducted according to the guidelines provided by the Cantonal Veterinary Office Basel-Stadt, and performed in compliance with the Swiss Veterinary Law.

Experiments were performed on mice from the following strains: C57BL/6JRj (Janvier), Fos^tm2.1(icre/ERT2)Luo^/J (TRAP-2) (Cat. No. 30323, Jackson Labs), and TRAP-2 mice crossed with B6.Cg-Gt(ROSA)26Sor^tm9(CAG-tdTomato)Hze^/J (Flex-tdTomato) (Cat. No. 7905, Jackson Labs). Mice were housed in temperature- and humidity-controlled cages, and kept on a 12h light-dark cycle with food and water provided ad libitum.

At the start of every experiment, mice were randomly assigned to an experimental group by the responsible researcher.

### Labelling isochronic neurons

To precisely monitor the developmental age of the embryos, 2 females and 1 male were put together in a freshly prepared cage and separated the following morning. At the moment of separation, the presence of a vaginal plug was considered a sign of effective copulation. Separation of the male from the females marked embryonic day 0 (E0). Labelling of isochronic neurons was performed using two strategies. For the first, we injected 5-bromo-2′-deoxyuridine (BrdU, 10 mg/ml in saline, 100 mg/kg, Sigma) through an intraperitoneal (*i.p.*) route on a specific embryonic day (from E11 to E16) in different cohorts of pregnant dams. BrdU is incorporated and permanently retained in the DNA of neurons undergoing cell division at the time of injection. Once labelled mice reached adulthood (older than 60 post-natal days, P60), the proportion of BrdU+ neurons within the CA3 network was visualized through histological means.

The second strategy aimed at achieving selective expression of Designer Receptors Selectively Activated by Designer Drugs (DREADDs) in hippocampal isochronic neurons. For this, we used an all-viral, two-step approach. The first step was performed during embryonic development, where an AAV vector for the expression of the recombinase Cre (pENN.AAV.CAMKII0.4.Cre.SV40, Cat. Number #105558-AAV1, titre 1.9 x 10^13^, Addgene) was injected *in utero* into the ventricle of the embryonic brain (as described in Donato et al. 2017, *67*) at E12 or E15, to target EBNs and LBNs, respectively. In the second step, a virus for the Cre-dependent expression of DREADDs (pAAV-hSyn-DIO-hM4D(Gi)-mCherry, Cat. Number #44362-AAV1, titre 2.3 x 10^13^; or pAAV-hSyn-DIO-hM3D(Gq)-mCherry, Cat Number #44361-AAV1, titre 2.2 x 10^13^, Addgene) was delivered locally to the dorsal (AP: -1.7 mm; ML: +/-1.8 mm; DV: -1.8, - 1.55 mm, and -1.3 mm, 200 nl injected at each location) and intermediate (AP: -2.3 mm; ML: +/-2.45; DV: -1.9 mm, -1.65 mm, and -1.4 mm, 200 nl injected at each location) hippocampus of Cre-expressing mice. Injections were performed through a stereotaxic frame (Kopf) using a Nanoject II / Nanoject III setup (Drummond) connected to a virus-loaded sharp glass capillary at a rate of 3 nl/s.

### Anaesthesia and analgesia during surgeries

On the day of the surgery, mice were injected subcutaneously (*s.c.*) with Buprenorphine (Bupaq P, 0.1 mg/kg, Streuli) and Atropine Sulphate (0.05 mg/kg, Amino AG) as pre-emptive analgesia, before being anaesthetized with isoflurane in oxygen (Piramal, 5% at induction, 1-2% for maintenance; 0.5 l/min airflow). Lidocaine (< 7 mg/kg, Streuli) and Bupivacaine (1-5 mg/kg, Sintetica) were administered locally through an *s.c.* injection at the beginning of the surgery. After the surgery, mice were allowed to recover under a heat lamp, and Buprenorphine was administered 4 to 6 hours after surgery. Meloxicam (Metacam, 5 mg/kg, Boehringer Ingelheim) was injected *s.c.* once a day for the following 3 days as postoperative analgesia.

### Viral injections at P2

For calcium imaging experiments, a viral vector for the expression of the genetically encoded calcium indicator GCaMP6f (pAAV.Syn.GCaMP6f.WPRE.SV40, Cat. Number #100837-AAV1, titre 1.3 x 10^13^, Addgene *95*) was injected stereotaxically (Kopf) into the hippocampus of the right hemisphere of mouse pups on postnatal day 2 (P2) (AP: +1.0; ML: +1.3; DV: -1.6mm). On the day of injection, pups were anaesthetized one by one with 5% isoflurane in oxygen (1 l/min airflow), delivered through a custom-designed nose cone (https://github.com/donatolab/isoMaskMousePups) within the frame of a stereotaxic surgery setup. The branching point where the medial and lateral sinuses connect over the cerebellum was used as a reference to calculate the location for the injection.

### Lens implants and base plating in adults

After stereotaxic alignment under isoflurane anaesthesia, the skull of adult mice was scored with a sharp needle tip to increase bone surface, and covered with a thin layer of Histoacryl (B. Braun Surgical). A 1.5 mm wide circular craniotomy was drilled, and the dura was removed before inserting an endoscope composed of a tubular, 1 mm wide GRIN lens glued to a 1 mm prism to gain optical access to the CA3 area of the hippocampus (custom needle endomicroscope design. Singlet tubular lens, diameter 1.0 mm, wavelength 520nm, non-coated. Prism: 1.00 x 1.00 x 1.00 mm, aluminium coating on hypotenuse. endomicroscope length approx. 4.67 mm (including prism). Object side: working distance 0.15 mm in water from prism exit surface / NA 0.4. Image side: working distance 0.08 mm in air / NA 0.5. GRINtech GmbH, Jena, Germany). The imaging face of the prism was implanted at a 45-degree angle facing the posterior-medial part of the brain, to follow the curvature of the hippocampus in the AP axis with its most frontal edge positioned at the following coordinates: AP: -1.58 mm, ML: +2.0 mm, DV: -2.3 mm. After insertion of the prism, the hippocampus was compressed by 0.1 mm towards the medial direction to secure stability and perfect adherence of the tissue to the imaging face of the prism. To secure the lens to the exposed skull, UV-curable cement (Venus Diamond Flow, Kultzer) was used at the interface between the lens and the skull, and secured with an additional layer of superglue (SuperGlue 415 Loctite, Loctite). Dental cement (Paladur, Kulzer) mixed with charcoal powder (Graphite, Carl Roth) was applied over the exposed skull and used to secure a metal head bar levelled with the surface of the endoscope. The exposed portion of the lens was protected with Kwik-Cast (World Precision Instruments).

1-2 weeks following lens implantation, a baseplate was permanently fixed to the head of the mouse to ensure easy mounting of a miniaturized, 1-photon microscope (Inscopix). For this, mice were kept under light isoflurane anaesthesia in a custom-made frame. The miniscope was attached to a baseplate, placed over the lens, and lowered to an optimal focal distance where active neurons were visible. The baseplate was fixed to the rest of the implant with UV-curable cement and secured with superglue. The miniscope was then removed, leaving the baseplate permanently fixed on the head of the mouse. Dental cement mixed with charcoal was applied around the UV-curable cement for extra protection and light shielding. A cap was placed on top of the baseplate to protect the lens in between imaging sessions.

### Fear conditioning tasks

Mice were trained in 4 variations of a fear conditioning task: (1) the contextual fear conditioning task; (2) the spaced contextual fear conditioning task; (3) the trace fear conditioning task; and (4) the second-order trace fear conditioning task.

During the Acquisition session of the *contextual Fear Conditioning* (cFC) task, mice were introduced into the fear conditioning setup (Hardware: Fear Conditioning System, Ugo Basile; Software: EthoVision XT, version 14-17, Noldus) and allowed to freely explore an environment (the Conditioning Stimulus, CS) consisting of a square box (Training Context, 25 x 25 cm floor area) characterized by distinctive visual patterns on the 4 walls, a specific odour (2% acetic acid), and an electrifiable grid floor. After 3 minutes, animals received a series of 5 foot shocks (Unconditioned Stimulus, US: 0.8 mA for 1 second) at 30 s intervals from each other. In the Recall session, performed at various intervals from Acquisition (from 1 up to 28 days), mice were re-exposed to the CS by placing them back into the Training Context, and allowed to freely explore the box for 5 minutes without the presentation of the US. As control, different cohorts of mice were either never experiencing the US during the Acquisition session (“Context only” control), or were not re-exposed to the CS on the Recall day (“No Recall” control), or were placed in a different context than the Training Context on a Recall day (Context B: circular, 27cm diameter, scented with 0.25% benzaldehyde in 70% Ethanol; “Context B Recall” control).

For the *spaced cFC* task, mice were subjected to two cFC Acquisition sessions, each of which involved the presentation of a lower number of US than the regular cFC task, and one Recall session. During the first acquisition session (Training), mice were placed in the Training Context (same as cFC task, CS) for 4 minutes before receiving 2 foot shocks (same as cFC, US) 30 seconds apart, with a total session length of 5 minutes. This learning episode was repeated during a second acquisition session (Re-Training), performed at an arbitrary temporal delay (Inter Session Interval, ISI), between 1 and 4 days from Training. During the Recall sessions, mice were reintroduced in the Training Context and allowed to freely explore for 5 minutes without the presentation of the US. For this experiment, two control groups were considered, both of which underwent only the Training and Recall sessions (*i.e.*: no Re-Training): one cohort experienced a low number of US presentations at Training (“2-US” Mass training regime) while another experienced a higher number of US at Training (“5-US” Mass training regime).

In the *trace Fear Conditioning* (tFC) task, during the Acquisition session, mice were introduced into the Training Context (same as the cFC task, Context-CS) and allowed to freely explore the environment for 3 minutes before being presented with a tone (Tone-CS, pure tone of 1 kHz, continuously for 20 s) that was followed, with a temporal delay of 30 seconds (Trace), by a foot shock (US; 0.8 mA for 2 s). The Tone-CS/Trace/US pairing was repeated 6 times, at various Inter-Trial-Intervals between repetitions (ITI = 120 s, 100 s, 140 s, 120 s, and 140 s). After the presentation of the last pairing, mice remained in the Training Context for 100 seconds before being shuttled back to their home cage. 1 or 28 days after Acquisition, mice were subjected to a Recall session. Different cohorts of animals were exposed to two types of Recall: in the “Context Recall” group, mice were placed back in the Training Context and allowed to freely explore for 5 minutes; in the “Tone Recall” group, mice were introduced into a new context (Context B, as in the cFC task), and allowed to freely explore the context for 3 minutes before being exposed to the Tone-CS 6 times (Tone-CS duration: 20 s. ITI = 60 s). Mice were then allowed to explore Context B for an additional 60 seconds before being shuttled back to their home cage.

Lastly, in the second-order *tFC* task, during the first-order Acquisition session (fo-Acq), mice were trained with the presentation of a first-order CS (a tone, as in the tFC task) followed, after a Trace delay (30 s), by a foot shock (US), as already described for the tFC Acquisition task. At an arbitrary Inter Session Interval (ISI) between 1 and 4 days from the fo-Acq session, mice were then subjected to a second-order Acquisition session (so-Acq), during which they were introduced into a novel context (Context B, as in the cFC task), and allowed to explore this context for 3 minutes. After that, mice were exposed to 6 pairings where a second-order CS (so-CS, White Noise, presented for 20 seconds) was immediately followed by the presentation of the first-order CS (fo-CS, tone, presented for 20 seconds), without the presentation of any US. The so-CS/fo-CS pairing presentation was repeated 6 times, with ITI = 40 s. In the Recall session on day 5, mice were introduced to Context B, allowed to explore the context for 3 minutes, and then presented 6 times with the so-CS (White Noise, 20 s duration, ITI = 60 s). For this experiment, control mice were only presented with the so-CS (white noise) without any presentation of the fo-SC (tone) during the so-Acq session (Unpaired Controls group).

During all experiments, mouse behaviour was recorded with an overhead infrared camera (Basler acA1300-60gm, Basler GenICam). Video recordings were processed offline, and memory retention and expression were quantified by the amount of time the animal spent freezing upon exposure to the CS during the Recall Sessions. Freezing bouts were systematically identified by the EthoVision XT software, and defined as moments of complete immobility apart from breathing (pixel change < 1-3% adjusted for each mouse, for at least 2 seconds). 60 to 90 minutes following the last session, mice were transcardially perfused as described below.

### Pharmacogenetic manipulations

For pharmacogenetic manipulation of EBNs/LBNs, a solution of clozapine-N-oxide (CNO (Sigma), 1 mg/Kg in 10% Dimethylsulfoxide (DMSO, Carl Roth), diluted to a final concentration of 1 mg/ml in NaCl), was injected *i.p.* 30 minutes before the time of intended maximum silencing/activation efficiency.

Throughout our chemogenetic manipulation experiments, and unless otherwise stated in the text, our Control groups included a pool of multiple conditions, among which: animals that received no viral nor CNO injection; mice that received a viral injection but no CNO injection; and animals that received no viral injection but a CNO injection. On histological and behavioural levels, animals belonging to these different conditions were strikingly similar in their levels of freezing or c-Fos expression, thus pooled together.

Mice in which the fraction of mCherry+ neurons was lower than the lowest observed fraction of BrdU+ neurons from pulse-chase experiments (*i.e.* 10%, Fig. S4B) were excluded from further analysis.

### Permanent labelling of Engram neurons

To permanently label Engram neurons, a solution of 4-Hydroxytamoxifen (4-OHT (MedChemExpress), 50mg/kg in 10% DMSO, diluted to a final concentration of 100mg/ml in sunflower seed oil (Sigma)) was injected *i.p.* immediately following the end of the cFC Acquisition session in TRAP-2/Flex-tdTomato mice. Quantification of tdTomato expression 1 or 2 days after 4-OHT injection was dependent on immunolabelling for the enhancement of tdTomato visualization. Thus, all brains with permanently labelled Engram neurons were stained for RFP (see Tissue processing and histology).

### Calcium imaging recordings

Time-lapse imaging of calcium activity in CA3 neurons from freely behaving mice was conducted using an integrated miniature fluorescence 1-photon microscope (nVoke 3.0 or nVue 1.0, Inscopix). A cohort of 5 mice was subjected to imaging during the Acquisition and Recall phases of the cFC task. Additionally, a second cohort of 5 mice was imaged while exploring the Training Context without the presentation of the US (similar to the “Context Only” control as previously described). Time-lapse recordings were longitudinally performed over 3 distinct imaging sessions, which included the day of Acquisition, 1 day and 14 days Recall.

At the start of each imaging session, the miniscope was affixed onto the baseplate while the animal had the freedom to run on a planar wheel while being head-fixed to a stable head-post. Following release from the holding setup, the animal was returned to its home cage, and CA3 network activity was recorded for 10 minutes before the behavioural training. This baseline period served as a normalization reference for spontaneous changes in the rate of calcium activity from individual neurons over consecutive days. Subsequently, mice were transferred to the Training Context without removing the miniscope, maximizing the likelihood of longitudinal tracking of the same neurons across experimental sessions. Recordings were paused during the handling and transfer periods. Calcium activity was recorded throughout the cFC or Context Only exploration sessions.

Recordings were obtained at rates of 20 or 25 Hz with a resolution of 1280 x 800 pixels, covering a Field of View (FOV) of 1000 x 625 µm (1 pixel = 0.765 µm^2^). On the first experimental day, imaging parameters such as the excitation LED brightness, gain, and focus plane were empirically optimized (Table S1) and maintained constant throughout the experiment. A histogram depicting the cumulative fluorescence intensity distribution for all pixels of the FOV was used to calibrate the LED intensity and maintain saturation rates between 12% - 75%. The focus plane was initially set to include the maximum number of neurons during the first imaging session and remained constant across sessions to maximize the longitudinal tracking of individual neurons.

### Processing and Analysis of Calcium Imaging Data

The raw recordings were processed using the Inscopix data analysis software Matlab API (IDAS, Version 1.6.0, Inscopix), as well as custom Matlab and Python scripts (Version 2021b, Mathworks).

As a pre-processing step, time-lapse recordings acquired on a specific day in the Home Cage and Training Context were concatenated and processed together. The recordings were spatially down-sampled by a factor of 2, resulting in a final FOV size of 640 x 400 pixels (1 pixel = 1.530 µm^2^). The FOVs from different imaging days were aligned to improve the registration of cells over time by matching landmarks such as blood vessels and lens corners from a transparent overlay image obtained from the FOV on the first day to the successive recordings.

To remove high and low spatial frequencies from the pre-processed data, pre-processed frames were filtered with a Gaussian blur band-pass filter with a high cut-off of 0.500/pixel and a low cut-off of 0.005/pixel. Motion registration was performed on a frame-by-frame basis using a previously published algorithm (*96*). For image segmentation, Constrained Nonnegative Matrix Factorization for micro-endoscopic data (CNMFe, provided by IDAS) was utilized to identify regions of interest (ROIs) corresponding to individual neurons, which served as the basis for subsequent analysis. Default values of IDAS were used for most parameters, except the minimal peak-to-noise ratio (15±4.4) and minimal pixel correlation (0.91±0.04), which were adjusted empirically for each animal. Cell-size parameters were derived empirically from recorded movies and set to 12 pixels diameter (18.36 µm^2^) after downsampling. ROIs were obtained by delineating a boundary that included only pixels with an average fluorescence value within the highest 20% range from the maximum pixel intensity for each ROI.

To extract calcium events from the raw calcium time series, custom Python scripts were developed. The raw calcium data were pre-processed by applying a 1^st^-order Butterworth low-pass filter (0.5 Hz cut-off) and compensating for any slow drifts or bleaching effects by fitting a polynomial model (order 1 or 2) to a low-pass filtered version of the signal (0.1 Hz cut-off). This model was subtracted from the low-pass filtered calcium data, effectively removing bleaching and other nonlinear trends in the calcium baselines. The standard deviation of the filtered calcium trace for each cell was computed. The 50^th^ percentile in the distribution of standard deviations was identified as the global minimum threshold for defining calcium events. This approach exploited data from all recorded neurons to establish a robust minimum value for the standard deviation, thereby minimizing the inclusion of spurious calcium events, particularly in low signal-to-noise (SNR) neurons. Calcium events were defined as periods when the fluorescence trace exceeded the global minimum (scaled by a constant). These events were further processed to distinguish the “up-phase” (representing the rising phase of the calcium event, determined by computing the derivative of the filtered calcium signal and selecting values > 0) and the “on-phase” (representing the entire period of the candidate event above the scaled global minimum).

To filter out false positives, only phases comprising continuously recorded frames with a minimum duration of 0.3 seconds for the up-phase events and 0.5 seconds for the on-phase events were considered. For each cell, SNR was calculated as the ratio of the mean event amplitude to the standard deviation of the noise (*97*). Noise statistics were obtained from the dF/F_0_ filtered calcium signal excluding up-phase events. To exclude calcium events from the noise statistics, we removed periods extending from 1 second before to 10 seconds after each up-phase event. If no up-phase events were present, the SNR was set to 0. The SNR thresholds for including cells in the analysed dataset were calculated based on the lowest 5-10% (±6.9±2.5) SNR values of each recording, and adjusted for each animal based on recording quality. Additionally, to account for instances where the algorithm over split traces associated with individual neurons, the pairwise distance between all ROIs was calculated, and if the distance was below a threshold of 5 pixels (half the width of an individual neuron), only the ROI with the higher SNR was retained for further analysis. Longitudinal tracking of neurons across sessions was conducted using dedicated algorithms provided by IDAS and based on previous work (*98*). A minimum correlation threshold of 0.6 was set to ensure reliable tracking between sessions. Subsequently, calcium traces associated with individual neurons were binarized: frames corresponding to the “up-phase” of calcium events were considered active and assigned a value of 1, while all other frames were assigned a value of 0. Only cells with an average activity rate of at least 0.01 Hz (i.e., active frames per second) in each of the 3 sessions were selected for further analysis.

To quantify the extent to which individual neurons modulated their activity rate in response to exposure to the Training Context compared to the Baseline period (*i.e.* in their Home Cage), we calculated a “Tuning Score” (TS) for each neuron and session (*74*). The TS was determined by computing the difference in activity rate (expressed in Hz) between the Training Context (TS) and the baseline activity rate in the Home Cage (HC), normalized by their sum:

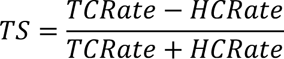

TS values across the population of recorded neurons were widely distributed, from neurons exclusively active in the Home Cage (TS = -1) to neurons exclusively active in the Training Context (TS = 1). Neurons with TS ≥ 0.33 (*i.e.* having double the activity rate in the Training Context than Home Cage) were classified as “Activated”; neurons with TS ≤ -0.33 (*i.e.* having half the activity rate in the Training Context than Home Cage) were classified as “Inhibited”; while neurons with -0.33 < TS < 0.33 were classified as “Neutral”. Values of TS for individual neurons across each pair of sessions (“Acquisition” vs. “Recall 1d”; “Acquisition” Vs “Recall 14d”; “Recall 1d” vs. “Recall 14d”) were plotted against each other and the underlying probability distributions were calculated using 2D-kernel density estimate (2D-KDE) to generate 2D density plots (custom-built scripts in Python 3.7 using *scipy* and *matplotlib*).

### Tissue processing and histology

On the last day of the experiment, mice were transcardially perfused with 4% PFA (Sigma Aldrich) in PBS. Before perfusion, deep anaesthesia was induced with isoflurane (5% in 0.5 l/min flow) and *i.p.* injection of Ketamine/Xylazine (Ketanarkon, 250 mg/kg, Streuli; Xylazine, 2.5 mg/kg, Streuli). Brains were extracted and kept at +4°C overnight in PFA, which was then replaced by a solution of 30% sucrose (Sigma Aldrich) in PBS.

Brains were frozen and cut into 50 µm sections on a cryostat, which were collected into 24-well plates. Sections for immunohistological staining were incubated in a blocking solution consisting of 10% Bovine Serum Albumin (BSA, Sigma-Aldrich) in PBST (1X PBS + 0.3% Triton-X100 (Roth)), for 1 h at room temperature under continuous agitation. After blocking, sections were incubated in a solution of PBST containing 3% BSA overnight at +4°C under continuous agitation, with one or a mix of the following primary antibodies: Rat anti-c-Fos (1:1000, Cat. No. 226 017, Synaptic Systems, RRID: AB_2864765), Rabbit anti-c-Fos (1:1000, Cat. No. 226 003, Synaptic Systems, RRID: AB_2231974), Guinea Pig anti-NeuN (1:1000, Cat. No. ABN90P, Sigma), and Rabbit anti-RFP (1:1000, Cat. No. 600-401-379, Life Technologies). Thereafter, sections were washed 3 times for 15 minutes each time at RT with PBST. Secondary antibodies (Chicken anti-rat 647 (1:500, Cat. No. A21472, Invitrogen), Goat anti-rabbit 647 (1:500, Cat. No. A21244, Invitrogen), Goat anti-guinea pig 488 or 548 (1:500, Cat. No. A11073/A11075, Invitrogen), and Donkey anti-rabbit 568 (1:500, Cat. No. A10042, Invitrogen)) incubation was performed for 2 h at room temperature under continuous agitation in PBST containing 3% BSA. Following this step, sections were washed once with PBST, and twice with 1X PBS, before being mounted on glass slides and protected by a coverslip using a custom-made anti-fade mounting medium (15%, Sekisui, Selvol 205; 30% Glycerole, 2.5% DABCO (Sigma, D27802) in PBS). Coverslip edges were sealed with nail polish 24h after mounting, and stored at +4°C in darkness until imaging.

Brains with BrdU labelling underwent an additional antigen-retrieval step to expose intercalated BrdU molecules. For this, after having stained for other markers, sections were incubated in a 1M solution of HCl (diluted from 25% HCl, Emsure) for 30 minutes at 45°C, then washed twice with 1X PBS, before being further incubated for 10 minutes at room temperature with a 1M solution of Borate buffer to neutralize HCl and terminate the antigen-retrieval protocol. Sections were then washed twice with PBST before being incubated with the primary antibody (Mouse anti-BrdU, 1:1000, Cat. No. B35128, Invitrogen; or Rat anti-BrdU, 1:1000, Cat. No. ab6326, Abcam, RRID: AB_305426) and the secondary antibody (Donkey anti-mouse 488 or 568 (1:500, Cat. No. A21202/A10037, Invitrogen) or Chicken anti-rat 488 (1:500, Cat. No. A21470, Invitrogen)) as previously described.

### Confocal imaging and histology quantifications

For image analysis, z-stacks were acquired using an Olympus Spinning Disk confocal microscope (Evident IXplore Spin Confocal Imaging Microscope System) equipped with Yokogawa CSU-W1 spinning disk unit and Hamamatsu ORCA-Fusion camera, Olympus). Different objectives were used: a 4x air objective (UPL X APO; NA: 0.16; single plane images), a 10x air objective (UPL X APO; NA: 0.4; z-stack step size = 2 µm), and a 30x silicone oil objective (UPL S APO; NA: 1.05; z-stack step size = 1 µm).

To ensure consistency in the quantification of variables such as c-Fos expression, c-Fos/BrdU co-localization, td-Tomato/BrdU/c-Fos co-localization, and mCherry expression, all images were acquired using identical imaging parameters. These parameters were determined at the beginning of the study, using an adult home cage control animal as a reference. The imaging parameters used for image acquisition were as follows: 488 channel: laser power = 30%, exposure time = 150 ms; 568 channel: laser power = 20%, exposure time = 100 ms; 647 channel: laser power = 30%, exposure time = 150 ms. To maintain consistency within each experiment, images from the same sample were acquired on the same day, and whenever possible, samples from the same experiment on consecutive days.

To perform a spatially unbiased analysis of positive cells distributed along the whole extent of the proximo-distal axis of the CA3 pyramidal layer in the dorsal and intermediate hippocampus, a systematic image acquisition approach was employed. For each slice, 3-6 images were acquired, ensuring coverage of the entire extent of the pyramidal layer in the area of interest. This process was repeated for the acquisition of 3-7 slices per animal, covering an anterior-posterior portion of the mouse hippocampus ranging from -1.58 to -2.3mm relative to Bregma. To create composite images of the same region, a custom script (10.5281/zenodo.7701150) was used to stitch together individual images. All subsequent analyses were performed using Imaris software (Imaris x64, version 9.3.1 to 9.9.1, Oxford Instruments).

Using the automatic spot detection function in Imaris, c-Fos+, BrdU+, NeuN+, tdTomato+, and mCherry+ neurons were identified within the 3D volume of the acquired image z-stacks. Distinct expected diameters for identifying positive cells were defined to account for the nuclear or cytosolic localization of the target protein. Specifically, an expected diameter of 6 µm was used for the automatic identification of c-Fos+ and BrdU+ nuclei, while an expected diameter of 7 µm was used for the identification of NeuN+, tdTomato+, and mCherry+ neurons. To eliminate false positives, a quality filter threshold of 20 arbitrary units, as suggested by the software, was applied. However, due to the spotted nature of the BrdU staining, this filtering procedure could not be applied for BrdU nuclei quantification. Therefore, false positives for BrdU+ nuclei were manually inspected and removed. Only cells with somata located within the CA3 pyramidal layer were considered for further analysis. To determine co-localization between two or more markers, the 3D overlap of positive spots was quantified. The number of cells positive for each marker was normalized to the total number of cells that could potentially express that specific marker in the region of interest, as specified on a case-by-case basis throughout the manuscript.

For the analysis of Engram reactivation, the probability that a neuron would be positive to all markers (*i.e.* tdTomato, c-Fos, and BrdU) by chance was calculated as the compound probability of each event happening independently from the others (calculated based on the average frequency in which positive cells for that specific marker were observed across the population of mice from similar experimental conditions):

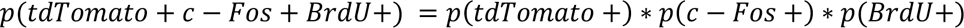

For our learning experiments, and unless otherwise stated in the manuscript, our control groups consisted of animals that were taken directly from their Home Cage, without exploration of the Training Context, at the time of tissue processing. Given the remarkable similarity in their baseline levels of expression of c-Fos in the CA3 network, Home Cage animals from different experiments were pooled and presented in multiple figures across the study.

To reduce potential bias in the analysis, the researcher conducting the analysis was blind to the experimental group to which animals belonged until after the data analysis was completed.

### Plotting and statistics

All Statistical analyses were conducted either on GraphPad Prism (GraphPad Prism version 9.5.1 and 10.0.0 for Windows, GraphPad Software, California) or Matlab (Matlab 2021b, Mathworks). For a specific description of the statistical analyses used throughout the study and precise reporting of *p* values and sample sizes, see Supplementary Table 2. Results were expressed as the mean ± standard deviation unless otherwise indicated. Individual samples were reported in each figure unless otherwise stated. *p* values (\**p*<0.05, \*\**p*<0.01, \*\*\**p*<0.001) and sample sizes are indicated in the legends, throughout the figures, and in Supplementary Table 2. Plots were originally produced in GraphPad, Matlab, or Python, and later modified using Illustrator (illustrator 2022, Adobe). No statistical tests were used to predetermine sample sizes, but our sample sizes were similar to those reported in previous publications.

